# Reciprocal host-*Wolbachia* interactions shape infection persistence upon loss of cytoplasmic incompatibility in haplodiploids

**DOI:** 10.1101/2024.11.18.624120

**Authors:** Felipe Kauai, Nicky Wybouw

## Abstract

Maternally transmitted symbionts such as *Wolbachia* spread within host populations by mediating reproductive phenotypes. Cytoplasmic incompatibility (CI) is a reproductive phenotype that interferes with embryonal development when infected males fertilize uninfected females. *Wolbachia*-based pest control relies on strong CI to suppress or replace pest populations. Host modifier systems have been identified that suppress CI strength, threatening the efficacy of *Wolbachia*-based pest control programs. In haplodiploids, CI embryos either die (Female Mortality, FM-CI) or develop into uninfected males (Male Development, MD-CI). The reciprocal spread of host suppressors and infection as well as the interaction with the two CI outcomes in haplodiploids remains poorly understood. The contribution of sex allocation distortion (Sd), an independent *Wolbachia*-mediated reproductive phenotype that causes a female-biased sex ratio, to infection persistency in haplodiploids is also poorly understood, especially with imperfect maternal transmission. To address these issues, we developed individual-based simulations grounded in the haplodiploid *Tetranychus urticae* pest system. We validated our simulations by tracking *Wolbachia* spread in experimental *T. urticae* populations and by contrasting the predictions of deterministic models. Within ∼15 host generations, we found that deterministic models inflate final infection frequencies by ∼8.1%, and significantly overestimate the driving potential of CI, particularly under low initial infection frequencies. Compared to MD-CI, we show that FM-CI strongly extends infection persistency when suppressors are segregating in the population. We also quantify how maternal transmission and CI outcome modulate the reciprocal spread of suppressors and infection. Upon loss of CI, we show that hypomorphic expression of Sd (∼5%) is sufficient for a stable persistence of infection. We derive a closed-form expression that approximates the stable polymorphic infection frequencies that can be maintained by Sd. Collectively, our results advance our understanding of how symbiosis with CI-inducing *Wolbachia* might evolve and inform CI-based pest control programs of potential future risks.

## Introduction

Maternally transmitted symbiotic bacteria, such as *Wolbachia*, commonly colonize the reproductive tissues of arthropods. These reproductive symbioses are typically not stable and infections are generally considered lost on a shorter timescale than host speciation and extinction (Atyame et al., 2011; Koehncke et al., 2009; Turelli et al., 2018, 2022). The persistency of infection is for instance lowered by imperfect maternal transmission and fitness penalties of infection (Hague et al., 2020, 2022; Hoffmann et al., 1990). Despite imperfect transmission and infection penalties, reproductive symbionts are able to spread within populations by altering host reproduction and increasing the proportion of infected transmitting females (Engelstädter & Hurst, 2009; Kaur et al., 2021). A variety of symbiont-induced reproductive phenotypes has been observed, with cytoplasmic incompatibility (CI) currently considered the most common. CI disrupts the embryonic development when an infected male fertilizes an uninfected female (Shropshire et al., 2020). Whereas embryos that suffer from CI exhibit higher mortality rates regardless of their sex in diploid arthropods, the phenotypic outcomes of CI are more complex in haplodiploid arthropods (Breeuwer, 1997; Perrot-Minnot et al., 2002; Vavre et al., 2001). Haplodiploidy is a mode of reproduction found in ∼15% of all arthropods, in which unfertilized eggs develop into haploid males, whereas fertilized eggs generate diploid females (de la Filia et al., 2015). After fertilization in haplodiploids, CI can cause embryonic and juvenile mortality in female offspring (Female Mortality CI, FM-CI). CI-induced defects can also cause fertilized eggs to develop into viable adult males instead of females (Male Development CI, MD-CI) (Nguyen et al., 2017; Perrot-Minnot et al., 2002; Vavre et al., 2001a). Different embryos from a single CI cross can suffer from either FM-CI or MD-CI, collectively determining total CI strength (Nguyen et al., 2017; Wybouw et al., 2022). When females and males carry a related symbiont, CI is rescued, favouring the maternal transmission and spread of the CI-inducing symbiont (Shropshire et al., 2020). In *Wolbachia*, CI factors A and B (*cifA* and *cifB*) genes control the induction and rescue of CI (J. F. Beckmann et al., 2017; LePage et al., 2017). *cifB* genes are required in infected males for CI induction, while *cifA* genes also play a contributing role in some systems, but not in all (“one-by-one” versus “two-by-one” models). Across all systems however, *cifA* genes are essential for CI rescue in infected females (Adams et al., 2021; J. Beckmann et al., 2023; Horard et al., 2022; Shropshire & Bordenstein, 2019).

When considering mutually compatible *Wolbachia* variants, theoretical work suggests that selection on *Wolbachia* within a specific host does not directly act on CI strength, consistent with the discovery of several pseudogenized *cif* genes and inferred CI dysfunction events (Martinez et al., 2021; Meany et al., 2019; Prout, 1994; Turelli, 1994). Instead, *Wolbachia* are expected to become mutualistic and increase the relative fitness of infected females in diploids and in haplodiploids (Egas et al., 2002; Prout, 1994; Turelli, 1994; Weeks et al., 2007). In support of these predictions, transitions from a deleterious to a mutualistic infection for female fecundity have been inferred in a number of host systems (Fry et al., 2004; Vavre et al., 1999; Weeks et al., 2007). In haplodiploid hosts, maternally transmitted symbionts also induce sex allocation distortion (Sd), a reproductive phenotype that reallocates the maternal investment of resources from male to female offspring. Here, regardless of the paternal infection state, offspring of infected females exhibit a more female-biased sex ratio (Katlav et al., 2022; Vala et al., 2003; Wybouw et al., 2023). Symbionts such as *Cardinium* and *Wolbachia* are able to simultaneously induce CI and Sd (Katlav et al., 2022; Wybouw et al., 2023). Previous theoretical work has suggested that Sd could strengthen the CI drive and ensure that symbionts spread within host populations from low initial infection frequencies (Egas et al., 2002; Wybouw et al., 2023). Whether Sd is sufficient for stable *Wolbachia* infections with imperfect transmission upon CI loss, and whether Sd might facilitate the evolution of a mutualistic infection remains to be investigated.

A body of theory indicates that selection in diploid and haplodiploid hosts favors the evolution of male-specific suppression systems that reduce CI strength (Egas et al., 2002; Koehncke et al., 2009; Turelli, 1994; Vala et al., 2002; Vavre et al., 2003). Empirical studies of *Wolbachia*-infected insect and mite hosts broadly support this prediction by showing that the host male genotype is a strong determinant of CI strength (Cooper et al., 2017; Reynolds & Hoffmann, 2002; Wybouw et al., 2022). Yet, compared to CI loss through pseudogenization of *cif* operons, our mechanistic understanding of CI loss through male suppression is limited. In contrast to the assumptions of a monogenic basis for host suppression of CI in previous theoretical analyses (Koehncke et al., 2009; Vala et al., 2002; Vavre et al., 2003), genetic work on the haplodiploid spider mite *Tetranychus urticae* indicates that male suppression of CI can be underpinned by a complex polygenic basis (Wybouw et al., 2022).

How the interaction and (co)evolution of host and symbiont traits determine the prevalence and persistency of infection remains poorly understood, especially in populations with changing demographic variables and overlapping generations. Understanding the reciprocal effects of host and symbiont traits on infection persistency at the population level is moreover vital for the development of durable *Wolbachia*-based pest control (Gong et al., 2020; Ross et al., 2019; Turelli, 1994). These biocontrol programs rely on strong CI to either suppress pest populations by periodically releasing infected males or to establish pathogen-blocking *Wolbachia* in disease vector populations.

In this study, we set out to investigate infection persistency upon loss of CI in haplodiploids. We studied the impact of *Wolbachia*-induced FM-CI and MD-CI on the reciprocal spread of host suppressors and *Wolbachia* infection and quantified the effects of Sd on the rate of infection loss. We studied these infection dynamics in theoretical and experimental populations of the haplodiploid model system *T. urticae*. We implemented experimentally derived data to parameterize and validate an individual-based model (IBM), whose predictive capacity was directly compared with previously developed deterministic models of *Wolbachia* spread in haplodiploids. Simulations allowed us to disentangle and quantify the individual effects of host and symbiont traits on infection dynamics on short (∼15 host generations) and long (>500 host generations) timescales. First, we measured how *Wolbachia* invasions lead to measurable demographic changes at the host population level and quantified the relative efficacy of MD-CI and FM-CI as drivers of infection. Additionally, we studied how host suppressors interact with *Wolbachia*-induced FM-CI and MD-CI and the consequences of these interactions for infection persistency. Transitions from a neutral to mutualistic symbiosis were investigated by numerically deriving the threshold fecundity increase to maintain infection upon CI loss under various levels of Sd. We further present analytical work on the impact of Sd and derive a closed-form expression that predicts the stable polymorphic infection frequency in a haplodiploid population upon CI loss.

## Methods

### Model outline

We consider an individual-based model consisting of an initial haplodiploid population of size *P*_0_ with overlapping generations, where individuals have an explicitly determined sex, age (measured in days), and *Wolbachia* infection status (infected or uninfected). The infection is maternally transmitted with probability 1 − *μ*, where *μ* denotes the imperfect transmission rate. Females are diploid and carry 2*L* biallelic (0 or 1) loci. Males are haploid and carry only *L* of these loci. Individuals reach maturity at age *M*, upon which females are considered adult and start to oviposit haploid eggs, whereas males are allowed to fertilize adult females. For a fertilized female, a fraction *λ* of the eggs receives haploid sperm of the fertilizing male and develops into female offspring. All remaining eggs (1 − *λ*) remain unfertilized and develop into males. Additionally, all individuals live to age *D*, which signals death by age, but might also be lost prematurely due to drift (see further). We adopted first-male sperm precedence for our IBM analyses, a common mating system found in *T. urticae* and other haplodiploid arthropods (Ablard et al., 2014; Rodrigues et al., 2020; Satoh et al., 2001). This entails that the sperm of the first male that mates with a virgin female will fertilize all eggs throughout the female lifespan. Consistent with the mating behavior of many haplodiploids, males were allowed to fertilize multiple females. Finally, we implemented *Wolbachia-*mediated FM-CI, MD-CI, and Sd in our IBM analyses. These reproductive phenotypes result in distinct outcomes from specific genetic crosses within our theoretical populations. In Figure 1 we present a diagram with an overview of the main components of the model.

**Figure 1.**
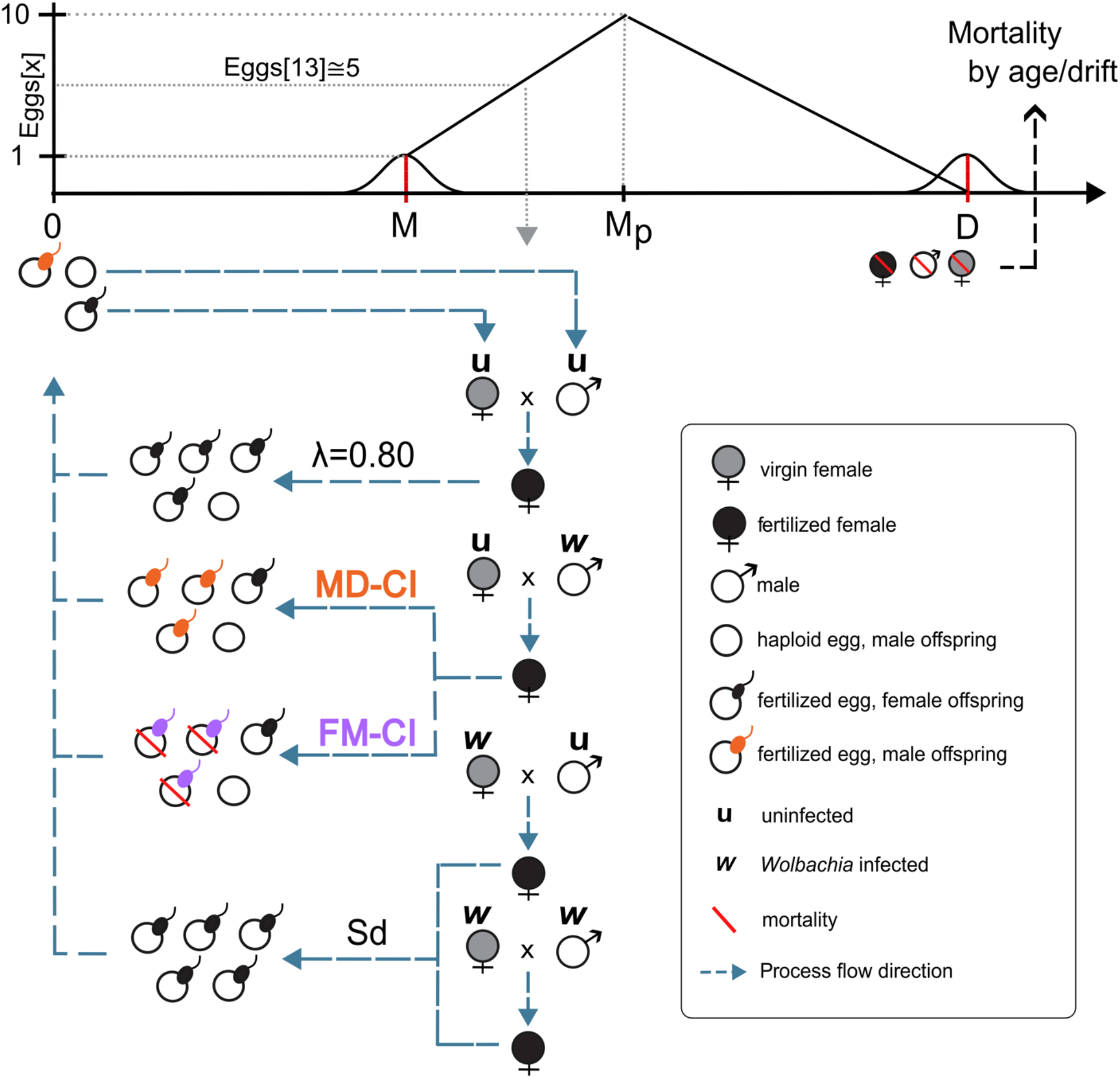
General outline of the individual-based model to study *Wolbachia* infection persistency in haplodiploid hosts. A hypothetical life cycle is depicted within the individual-based simulation, where the upper part represents the time line from oviposition (age 0) to death (age *D*), and a schematic representation of the expected number of eggs as a function of female age *x* (*Eggs*[*x*]). The number of eggs grows linearly from maturity (age *M*) to a maturity peak *M*_0_, after which it steadily decays. Individuals between age 0 and age *M* are considered non-reproducing juveniles. When individuals reach the maturity threshold, females and males are allowed to oviposit and fertilize eggs, respectively. We depict the four cross types dependent on the infection status of females and males (infected or uninfected). The reproductive outcomes are illustrated of a compatible cross, two incompatible CI crosses inducing either MD-CI or FM-CI, and the two Sd crosses. The illustrative cross types are exemplified using an adult female at age 13 laying 5 eggs. The flow of processes in the simulation is depicted by dashed blue lines.

### Haplodiploid reproduction and population growth

The expected daily oviposition (i.e., number of eggs per day) of an adult female is a function of its current age x, which we denote by Eggs[x]. The actual number of eggs that a female oviposits at a certain day is derived from a normal distribution, centered at Eggs[x] with standard deviation 0.5, and rounded to the closest integer. At the female maturity threshold M, we define Eggs[M] = 1 (the expected number of eggs laid by a female at age M is equal to one). Daily oviposition increases linearly as a function of female age until the maturity peak M_p_, after which daily oviposition linearly decreases until the female dies naturally at age D according to the following piecewise linear functions:

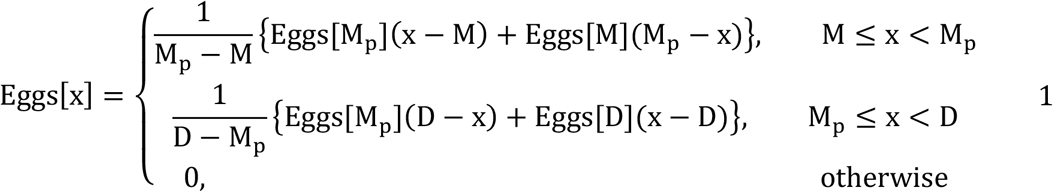

No linkage is assumed during our simulation of oogenesis. The set of loci (*L*) a haploid oocyte receives is therefore the product of a random selection of the maternal alleles. If the female is fertilized, then a proportion *λ* of all oocytes receives the male chromosome, resulting in diploid female offspring (2*L*). A proportion of the eggs equal to 1 − *λ* remains unfertilized and develops into haploid males.

Each iteration of our IBM corresponds to a day. For each day, the system recovers all females and males in reproductive age. If a female is virgin, then a male partner is chosen at random for mating, whereby the male loci are stored as an attribute of the female traits (i.e., the female is fertilized for life, following first-male sperm precedence). We determine the number of eggs that will be laid by the adult female following Equations 1, and repeat this procedure for all females in reproductive age in a given day. As the expected number of eggs for each female is greater than one and generations are overlapping, population sizes are expected to increase exponentially. To introduce an upper bound to the size of our haplodiploid populations, we simulated mortality that randomly targets juvenile and adult mites. This random mortality can be interpreted as the outcome of various biotic or abiotic processes (including predation and resource shortage). We constrained population growth to a logistic form with carrying capacity *K*. Let *P*(*t*) be the total population size at day *t* (the sum of eggs, juveniles and adults), then we compute the proportion *ψ*(*t*) of individuals that will die at the end of each day *t* as:

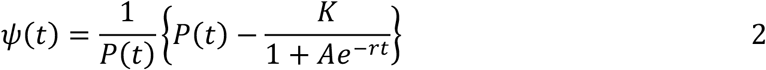

where *r* is the growth rate, and

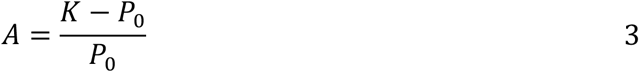

This implementation ensures that the total number of individuals grows logistically to the stable equilibrium *K*. Since *P*(*t*) grows exponentially with rate given by the expected number of eggs in Equation 1 (i.e., quicker than the logistic form), Equation 2 is well defined for all time *t* (see also further below). Importantly, because a proportion *ψ*(*t*) of the total population size dies at the end of any given day, irrespective of age, sex, or infection status, the population sex ratio is not affected, but is maintained by the proportion of fertilized eggs (controlled by *λ*). The remaining mites (with age less than *D*) survive to the next day and the process is repeated.

### *Wolbachia*-mediated reproductive phenotypes in a haplodiploid host

In our simulations, *Wolbachia* symbionts carry a *cif* gene repertoire that is characterized by three reference parameters. These parameters are real numbers in the closed interval [0, 1] and represent the CI phenotypes, FM-CI (*σ*_*fm*_) and MD-CI (*σ*_*md*_), as well as the total CI strength (*σ*_*total*_). The first two parameters, *σ*_*fm*_ and *σ*_*md*_, indicate the relative contribution of each CI phenotype to total CI strength. For instance, if *σ*_*fm*_ = 0.9 and *σ*_*md*_ = 0.1, an incompatible CI cross will induce an expected mortality of 90% of the fertilized eggs, with 10% of the remaining fertilized eggs developing into adult males with probability 0.1. To streamline our study, we assume that *σ*_*fm*_ + *σ*_*md*_ = *σ*_*total*_ in all simulations and that the relative proportions of the two CI phenotypes remain constant as we vary *σ*_*total*_. For instance, if we change *σ*_*total*_ from 1 to 0.5 and originally *σ*_*fm*_ = 0.9, then the new strength of the CI phenotypes will be *σ*_*fm*_^′^ ≔ *σ*_*total*_*σ*_*fm*_ = 0.45 and *σ*_*md*_^′^ ≔ *σ*_*total*_*σ*_*md*_ = 0.05.

In our model, *σ*_*fm*_ controls the hatching probability *θ* of fertilized eggs. We define the probability that juveniles emerge from uninfected eggs fertilized by infected sperm to be *θ* ≔ 1 − *σ*_*fm*_. We define *σ*_*md*_ as the probability that a fertilized egg develops into a functional male (instead of a female) that is composed solely by the maternal genetic information. Naturally, when *σ*_*md*_ = 1 and *σ*_*total*_ = 1, all fertilized eggs from an incompatible CI cross develop into adult males. Note that first-male sperm precedence ensures that all eggs of uninfected females carrying infected sperm will be subjected to FM-CI or MD-CI, or both, throughout the female’s life span.

In crosses with infected females, offspring sex ratio can be altered by sex allocation distortion (Sd, Figure 1). In Sd crosses, after a proportion of the eggs has been fertilized according to *λ*, each of the remaining unfertilized eggs have a probability *α* of also becoming fertilized, thus producing a bias toward a higher proportion of female offspring. Note that Sd is a population level measure in the experimental literature (Katlav et al., 2022; Wybouw et al., 2023), and the probability of an additional egg being fertilized due to Sd is an individual property in our IBM. To couple Sd to this probability, suppose an infected female oviposits *N* eggs on a certain day. In the absence of Sd the expected number of females and males is *λN* and (1 − *λ*)*N*, respectively. The expected number of additional fertilized eggs due to Sd from a batch of unfertilized eggs can be computed as (1 − *λ*)*Nα*. Sd satisfies the following relation:

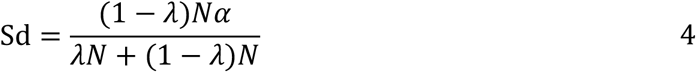

which implies,

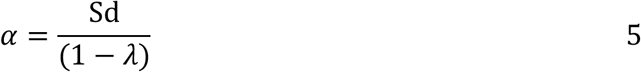

Moreover, we consider a possible fecundity increase (*F*) in infected females, a common feature of mutualistic *Wolbachia* infections (Hoffmann et al., 1990; Weeks et al., 2007). Similar to *α, F* denotes the probability that for each oviposited egg, an additional one will be produced. For instance, if *F* = 0.10, the expected number of additional eggs laid by an infected female will be 10% of the original batch given by Equations 1.

### Polygenic host modulation of *Wolbachia*-induced CI

In *T. urticae*, host modulation appears to be mainly mediated by infected males and behaves as a polygenic trait (Wybouw et al., 2022). To simulate host suppression, CI strength is modified by the chromosomal composition of the male genome. Each of the *L* loci represents a single CI suppressor and has two alleles; allele 0 codes for a non-active suppressor, whereas allele 1 codes for an active system. In the current study, we assumed that the host suppressors do not display epistatic interactions and that their effects on CI strength are strictly additive. Thus, under a strict FM-CI regime, the activation of each suppressor increases *θ* following an incompatible CI cross by an amount *σ*_*fm*_/*L*. For example, let *L* = 5 and consider an infected male with two active suppressors. Then, for *σ*_*fm*_ = 1, this infected male will suppress 2/5 of FM-CI, resulting in a hatch rate of 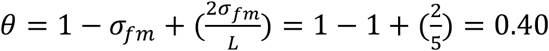. In this incompatible CI cross, each fertilized egg has a 40% hatching probability. The same principle applies for the control of MD-CI, by modulating *σ*_*md*_, thereby changing the probability that a fertilized egg develops into a haploid male. Note that, unlike the variation of CI through *σ*_*total*_, host suppression of CI does not necessarily maintain the relative proportions of *σ*_*fm*_ and *σ*_*md*_. Indeed, host suppressors may act either on *σ*_*fm*_, *σ*_*md*_ or both.

### *Wolbachia* transmission and spread within experimental *T. urticae* populations

The near-isogenic Beis-*w* and Beis-c lines were selected for the current study. Beis-*w* carries a single CI-inducing *Wolbachia* variant, whereas Beis-c is cured from infection by antibiotic treatment (Wybouw et al., 2022). Before the onset of the experiments, both lines were propagated for over ten generations on detached bean leaves (*Phaseolus vulgaris* L. cv ‘Prelude’) at 24°C, 60% RH, and a 16:8 light:dark photoperiod. We selected the Beis genotype because CI is complete against this host nuclear background. Prior to experimentation, the infection states of *Wolbachia* and other symbionts were verified by end-point diagnostic PCR assays, as previously described (Wybouw et al., 2022).

To study the maternal transmission efficiency of *Wolbachia*, age-synchronized cohorts of Beis-*w* and Beis-c were generated by a 24 hours oviposition period of 50 adult females on a detached bean leaf. During offspring development, 15 Beis-*w* females were isolated as teleiochrysalids (the last molting stage before adulthood) on a single day and were grouped with 8 one-to three-day old Beis-c adult males on a detached bean leaf. Beis-*w* females were given three days to emerge as adults and to mate with Beis-c males. Ten fertilized Beis-*w* females were placed on individual 16 cm^2^ leaf discs and were allowed to oviposit for 48 hours. Females were removed and offspring were individually collected upon reaching the deutonymphal stage. DNA samples were extracted from a total of 146 mites and were tested for infection using diagnostic PCR (Wybouw et al., 2022). DNA quality was verified for those mite samples that tested negative for *Wolbachia* by amplifying a fragment of the mitochondrial *COI* gene (Wybouw et al., 2022).

To study the spread of *Wolbachia*, age-synchronized cohorts of Beis-*w* and Beis-c were established by allowing 120 adult females to oviposit for 24 hours. For each experimental population, ∼30 females were isolated on a single day as teleiochrysalids and were paired with ∼20 one-to three-day old adult males of their own line on a detached leaf. Three days were given to the females to emerge as virgin adults and for fertilization to occur. The experimental populations were established on single potted bean plants by pooling 25 infected (Beis-*w*) and 25 cured (Beis-c) fertilized females in three batches of three replicates. New bean plants were offered to the nine expanding populations on a weekly basis. At 13 different timepoints (batch 1; at day 22, 47, 64, 75, 88, batch 2: at day 24, 51, 69, 81, 98, and batch 3: at day 25, 55, 99), 25 to 30 adult females were randomly isolated from each experimental population and were tested for *Wolbachia* infection (Wybouw et al., 2022).

### Model parameterization and analyses

In our model, we assumed that the number of eggs laid by a female grows linearly from an age signaling maturity (M = 11) up to what we refer to as a maturity peak (M_p_ = 16), where the maximum expected number of produced eggs is set to be *Eggs*[M_p_] = 10. Further reflecting the reproductive biology of *Tetranychus* mites, males are allowed to fertilize females upon reaching maturity, which we defined to be one day earlier than females (i.e., 10 days). All mites are allowed to live up to a maximum of 20 days. We allowed for individual variation within the populations by drawing parameter values from a normal distribution centered at the parameter average with a standard deviation of 0.5. These definitions clearly influence population growth, which we constrained by setting the carrying capacity to *K* = 5000 and computing *r* in such a way that by day 60 the population size reaches *K*/2. These decisions were made based primarily on computational constraints, since arbitrarily large values of *K* would render the model computationally intractable. For all analyses, unless stated otherwise, we used an initial population size of 50 (*P*_0_) adult females following our experimental setup. Sensitivity analyses revealed that these decisions had no effect on our results (see Results).

A single simulation consists of running the system for a total of 150 days (∼host generations), which we refer to as a reproductive year, and approximates the time interval in which we can observe active reproduction within natural populations of *T. urticae* in West-Europe (Helle & Sabelis, 1985; Xue et al., 2023). However, to understand the dynamics of reproductive symbiosis over longer time periods, we also ran simulations for multiple years, selecting a random subset of the population at the end of a year which served as the initial population for the next reproductive year. This process adds an additional drift component, as discussed within the next section. All results are based on a total of 100 independent runs of the simulations.

We set each egg to be fertilized with probability *λ* = 0.8 following recent experimental work in *T. urticae* (Wybouw et al., 2023). From a brood of eggs, we expect that 80% of the offspring will be female. The natural male to female ratio in the population should then remain at a proportion of 1:4. Indeed, in Supplementary Figure S1 we present the general descriptive statistics of the demographic features of the model, where the male to female ratio is normally distributed in the population by the end of a year with average 0.2501 and standard deviation 0.0063. In the model, 62.9% of all individuals die between the age of 0 and 3 days. Approximately 6.2% and 1.6% of females and males, respectively, successfully reach adulthood and reproduce. In accordance with our simulations (Supplementary Figure S2), copulation typically occurs as soon as females molt into virgin adults in natural population (Potter et al., 1976). In Table 1 we provide a summary of all parameters used in our model with their corresponding values and ranges. Additional technical information on model implementation can be found in Supplementary Material Text S1. Here, we discuss the logical structures of the simulations to facilitate reproducibility.

**Table 1.** Description of model parameters, with their respective symbols and values. For values taken from a normal distribution, their standard deviation is given in parenthesis. The studied ranges of values for parameters are highlighted by square brackets. For sex-specific parameters, we discriminate between males and females. When the parameter is independent of sex, we provide the corresponding value under the category *“general”*.

| Description | Symbol | Value |  |  |
| --- | --- | --- | --- | --- |
|  |  | Female | Male | General |
| Number of days in a reproductive season | $T$ | - | - | 150 |
| Total host population size at time $t$ | $P(t)$ | - | - | - |
| Carrying capacity | $K$ | - | - | 5000 |
| Proportion of individuals that will die at the end of a day | $\psi(t)$ | - | - | - |
| Age at which individuals die naturally | $D$ | 20 (0.5) | 20 (0.5) | - |
| Age at which individuals become mature | $M$ | 11 (0.5) | 10 (0.5) | - |
| Age at which females reach the highest daily oviposition rate (Maturity peak) | $M_p$ | 16 (0.5) | - | - |
| Expected number of eggs at maturity | $E[M]$ | 1 (0.5) | - | - |
| Maximum number of expected eggs at maturity peak $M_p$ | $E[M_p]$ | 10 (0.5) | - | - |
| Proportion of fertilized eggs, giving rise to female offspring | $\lambda$ | 0.8 | - | - |
| Imperfect transmission rate of *Wolbachia* | $\mu$ | [0, 0.05] | - | - |
| Number of loci in the host | $L$ | $2 \times 5$ | 5 | - |
| Female Mortality CI | $\sigma_{fm}$ | [0, 1] | - | - |
| Male development CI | $\sigma_{md}$ | [0, 1] | - | - |
| Total CI strength | $\sigma_{total}$ | [0, 1] | - | - |
| Increased relative fecundity in infected females | $F$ | [0, 0.10] | - | - |
| Sex allocation distortion | Sd | 0, 3 or 5% | - | - |
| Hatching probability | $\theta$ | [0, 1] | - | - |
| Initial *Wolbachia* infection frequency | $\omega_0$ | [0, 1] | - | - |

## Results

### The spread of *Wolbachia* in experimental and simulated Beis populations

Based on experimental work on the *T. urticae* Beis genotype (Wybouw et al., 2022), we parametrized our model using the observed strength of the two CI phenotypes (*σ*_*fm*_ = 0.98 and *σ*_*md*_ = 0.02). Maternal transmission of *Wolbachia* was imperfect in Beis mites and was estimated to be *μ* ≅ 1.8% (Supplemental Data). To assess the accuracy of our individual-based simulations of *Wolbachia* spread, we contrasted these values with the observed *Wolbachia* spread within our experimental populations and those predicted by deterministic models. Let *f*_*t*_ and *m*_*t*_ denote the frequency of infection among females and males at time *t*, respectively. Infection spread in haplodiploid populations can then be modelled by a set of non-linear coupled recursions (Egas et al., 2002; Vavre et al., 2000; Wybouw et al., 2023) as follows:

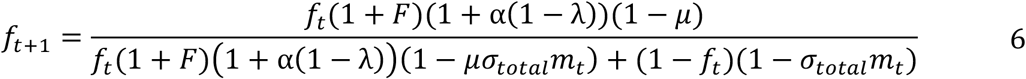

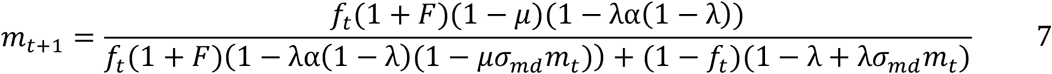

Using replicated experimental Beis populations with a 50% initial female infection frequency (*ω*_0_ = 0.5 in the IBM and *f*_0_ = 0.5 in the deterministic model), we observed a steady increase in *Wolbachia* infection frequency, reaching approximately ∼90% within a 100-day interval. The observed infection frequencies closely matched our simulations throughout the invasion process (Figure 2A), where the predicted final infection frequencies by the IBM were also ∼90% within the same time interval (the relative root mean squared error compared with experimental data was ∼17.54% (Figure 2A)). Equations 6 − 7 (with a generation time *t* of 11 days, following *T. urticae* biology) overestimated infection frequencies throughout *Wolbachia* spread and predicted an infection frequency of ∼98% by the end of 100 days. To test whether unparametrized constants had any appreciable effect on infection dynamics in the IBM, we also performed simulations with varying initial population sizes (*P*_0_) and carrying capacities (*K*). Although higher initial population sizes decreased the variance of the average daily infection frequency, we found that these quantities did not influence the rate and general pattern of *Wolbachia* spread (Supplementary Figure S3).

**Figure 2.**
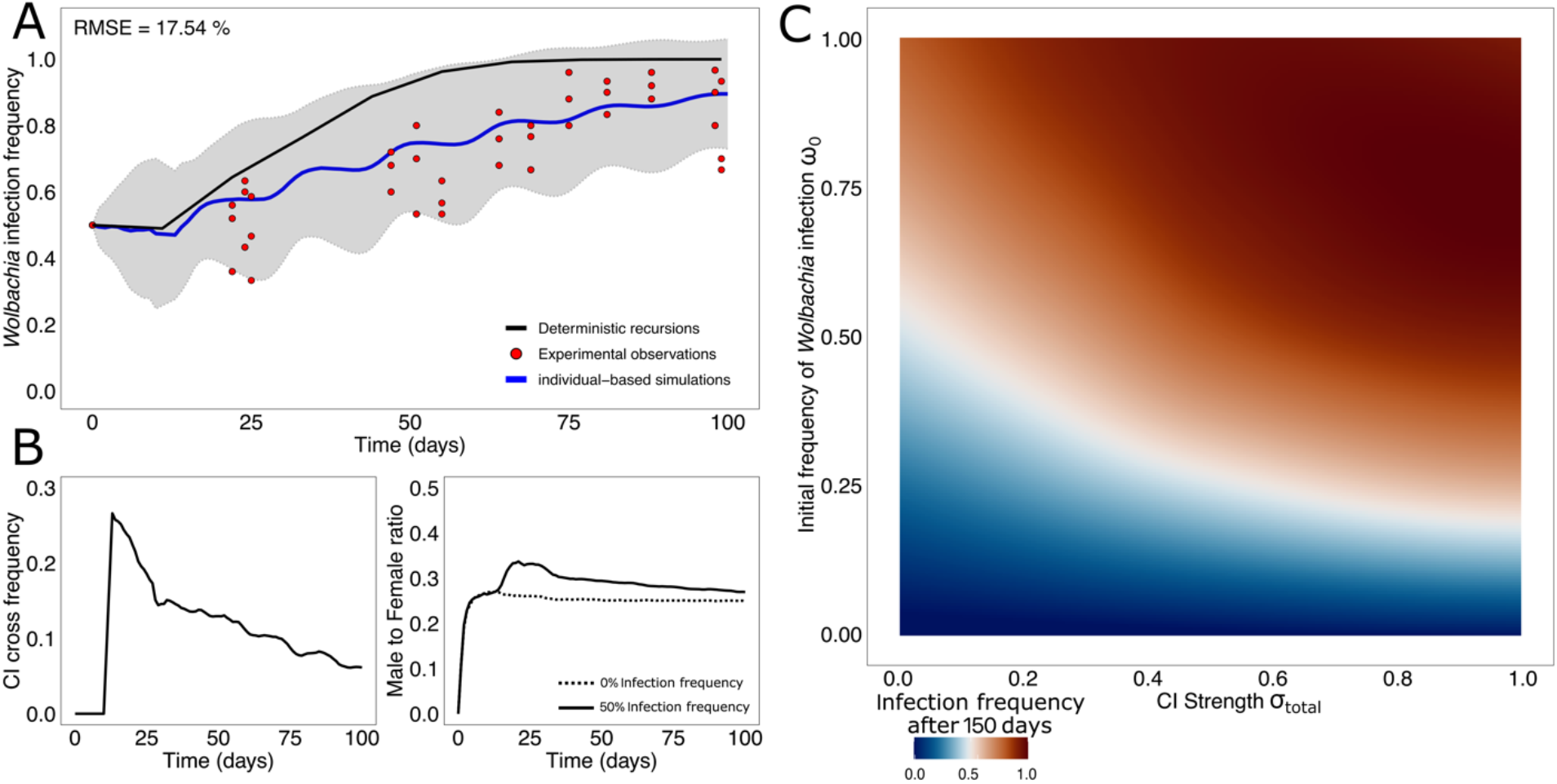
CI-inducing *Wolbachia* spread similarly in simulated and experimental Beis populations. For panel A, the predicted *Wolbachia* infection frequency by the IBM is plotted as a solid blue line as a function of time, with a shaded ribbon representing ± 1 standard deviation from the mean. The predicted infection spread by non-linear recursions are plotted as solid black line. Our observed *Wolbachia* infection frequencies are plotted as red dots (nine replicated Beis populations across three batches). Panel B depicts the frequency of CI crosses and male to female ratio for each day. Panel C depicts the final infection frequency at the end of single year (after 150 days of reproduction) across all values of CI strength (*σ*_*total*_ ∈ [0, 1]) and initial infection frequencies (*ω*_0_ ∈ [0, 1]). We implemented the empirically derived 2/98 MD-CI/FM-CI ratio for Beis mites. Final infection frequency values have been interpolated with a second-degree polynomial for enhanced visualization.

In our parametrized simulations of *Wolbachia* spread in Beis populations, the frequency of CI crosses peaked at day 15 (25.5%) which correlated with a first expressive increase of infection. This peak coincided with the maturity peak of oviposition (*M*_*p*_ = 15 and *Eggs*[*M*_*p*_] = 10), where F1 offspring generated at the start of the simulation is expected to reach maximum reproductive output. The frequency of CI crosses during *Wolbachia* spread also led to a disruption of the distribution of population sex ratios in time. Disrupted sex ratios reached a maximum average value of 33.2%, compared to the natural sex ratio that is centered at 25% (*λ* = 0.8) (Figure 2B).

These temporal changes in sex ratio are a consequence of haplodiploidy, since CI either kills females, or converts fertilized eggs into males (FM-CI and MD-CI, respectively). In either case, the relative frequency of males is expected to increase. These sex ratio disturbances were transient and only prominent during the early stages of *Wolbachia* spread, gradually decreasing together with the frequency of CI crosses. When *Wolbachia* infection approached fixation, CI crosses were halted and sex ratios returned to the natural state.

In our IBM, without CI, Sd, or fecundity effects (negative nor positive), initial infection frequencies need to be sufficiently high to overcome imperfect maternal transmission and stochasticity. As *σ*_*total*_ increases, the initial infection threshold frequency to obtain an expected final infection frequency of ∼50% by the end of a reproductive year (150 days) decreases non-linearly down to ∼20% when *σ*_*total*_ = 1 in our simulations (Figure 2C). *Wolbachia* are rapidly lost for initial infection frequencies under ∼10% independent of *σ*_*total*_. Importantly, we found that the deterministic recursions given by Equations 6-7 overestimate the ability of CI to drive infection relative to the IBM by an average of 8.1%. The difference is especially pronounced in the regime of high CI and low initial infection frequencies (Figure S4).

### Relative drive efficiency of the two CI outcomes and the transient impact on population sex ratio

In Beis populations, the relative contribution of MD-CI to total CI strength was ∼2%. However, the relative contribution of MD-CI in natural systems covers a continuum, with 0 and 100% as the extremes (Vala et al., 2002; Wybouw et al., 2022). To expand our understanding of the effects of these CI outcomes on *Wolbachia* spread and the host population, we analyzed the response of the model either under a strict female mortality (*σ*_*fm*_ ∈ [0,1] *and σ*_*md*_ = 0) or male development (*σ*_*md*_ ∈ [0,1] *and σ*_*fm*_ = 0) regime. Although no clear distinction in the final infection profile in function of CI strength and initial infection was found between FM-CI and MD-CI (Supplementary Figure S5), we did observe quantitative differences between these regimes. Contrasting the two CI regimes, we found that MD-CI decreased final infection frequency up to ∼12% when compared to FM-CI (Figure 3A). This observation is consistent with previous theory using deterministic models predicting that FM-CI facilitates symbiont invasion relative to MD-CI (Vavre et al., 2000, 2003). When initial infection frequencies were higher than ∼75% or total CI strength was lower than ∼40%, no substantial differences were observed between the two CI phenotypes. To further dissect the differences between MD-CI and FM-CI, we measured the frequency of CI crosses per day at the point of maximum divergence between the two CI phenotypes, i.e., for *ω*_0_ = 0.25 and *σ*_*total*_ = 1.0. We verified that, under female mortality, the frequency of CI crosses is consistently higher throughout the spread of infection (Figure 3B). This is a consequence of an overproduction of uninfected males by MD-CI relative to FM-CI, reducing the frequency of CI crosses that involve infected males. This transient overproduction of uninfected males can be clearly detected in population sex ratio, which peaked at ∼55% under a MD-CI regime compared to ∼33% for a FM-CI-inducing *Wolbachia* (Figure 3C).

**Figure 3.**
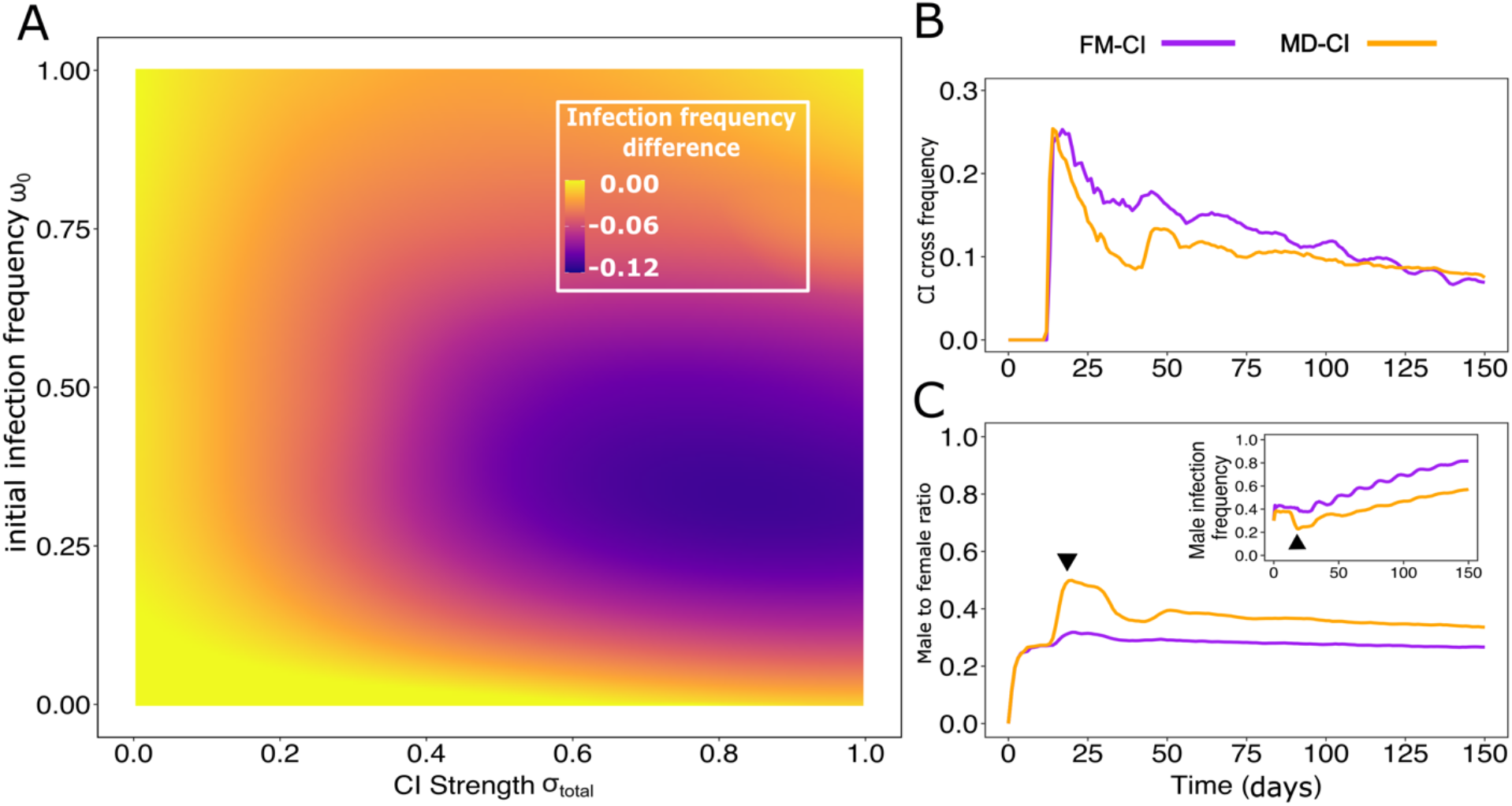
MD-CI constrains the spread of *Wolbachia* that induce strong CI by generating a transient higher proportion of uninfected males in a haplodiploid population. Panel A depicts the difference of the infection profile of a haplodiploid population under an MD-CI regime relative to an FM-CI regime. A marked difference of ∼12% is verified for *ω*_0_ ≈ 0.25 and *σ*_*total*_ = 1.0. Panel B depicts the frequency of CI crosses for a population with *ω*_0_ = 0.25 and *σ*_*total*_ = 1.0, under FM-CI and MD-CI. Under the same parametrization, panel C depicts population sex ratios (male to female) through time under FM-CI and MD-CI. The inset depicts the corresponding male infection frequencies. Black arrows highlight the correspondence in time of maximum male to female ratio with the minimum infection frequency among male hosts.

### Reciprocal effects of segregating host suppressors and CI-inducing *Wolbachia*

Host suppression is predicted to be a strong determinant of the spread and persistence of CI-inducing *Wolbachia* (Koehncke et al., 2009; Røed & Engelstädter, 2022; Turelli, 1994; Wybouw et al., 2022). Yet, our understanding of the reciprocal impact of host suppression and CI induction on the frequencies of *Wolbachia* infection and suppressor alleles remains limited. Here, we studied how polygenic suppression spreads through haplodiploid populations, and how these allele frequency changes impact *Wolbachia* prevalence and persistence.

We first initialized our model with 50% of randomly chosen host loci carrying active suppressors. Simulations ran for a period of 100 years (with one year defined to be 365 days) with ω_0_ = 0.50 under a strict FM-CI regime (*σ*_*fm*_ = 1, *σ*_*total*_ = 1), MD-CI regime (*σ*_*md*_ = 1, *σ*_*total*_ = 1) or upon loss of CI, simulating *cif* pseudogenization (*σ*_*total*_ = 0). At the transition of each year, we kept 100% of the population in the previous year as the new founding population, and therefore, *P*_0_ = *K* at all times after the simulation of the first year. By setting the population size at carrying capacity *K* through the transition of the years we erased the effect of drift, which is amplified when the population size is periodically reduced to *P*_0_ (see further).

In our CI dysfunction regime (simulating *cif* pseudogenization), the frequency of suppressors remained centered at 50% (Figure 4A). In contrast, we found that both FM-CI and MD-CI regimes invariably led to a fixation of suppressors (Figure 4A). These results are consistent with previous work that assumed a constant *Wolbachia* infection frequency and a monogenic basis of suppression (Vavre et al., 2001). Here, under MD-CI, we verified that the spread of suppressors was approximately twice as fast as under FM-CI, reaching near-fixation after ∼50 years. Since CI strength is gradually weakened, the spread of host suppressors coincided with a dramatic decrease in *Wolbachia* frequency under both CI regimes (Figure 4B). Note that we only identified expressive changes in the frequency of suppressors during long time frames, with negligible signals of change in allele frequency during a single year (∼35 host generations) (Figure 4C). Additionally, we found that Sd does not interact with the spread of host suppressors (Supplementary Figure S6).

**Figure 4.**
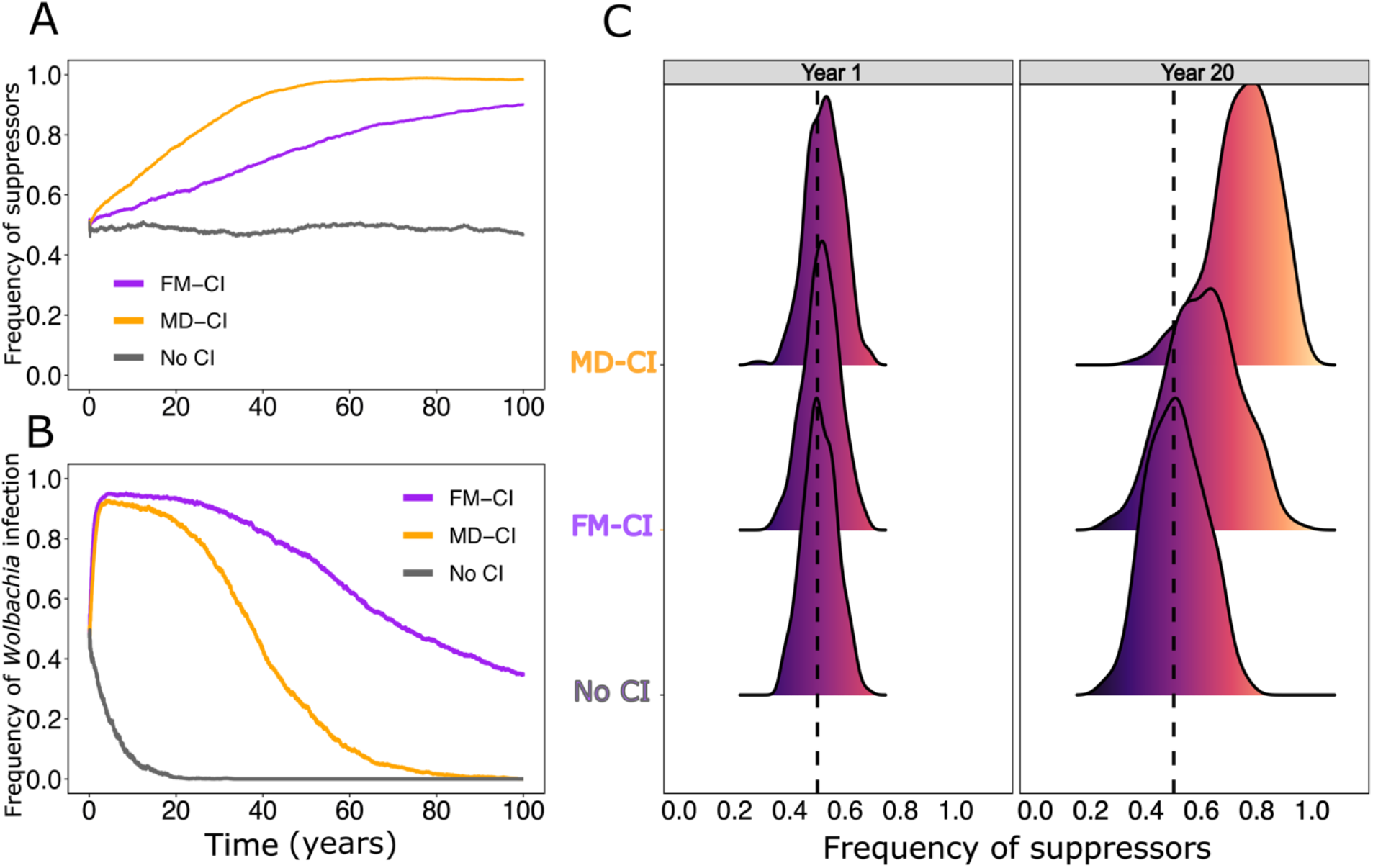
Host suppressors spread to near-fixation at different rates under FM-CI and MD-CI regimes. Panels A and B depict the temporal changes in the frequency of suppressors and *Wolbachia* infection, respectively. The dynamics with *Wolbachia* variants that induce FM-CI, MD-CI, and no CI are outlined. Panel C shows the frequency distribution of suppressors at year 1 and 20, factored by CI type (FM-CI, MD-CI, and no CI). In Figure S7, a diagram is presented to visually explain why an incompatible CI cross promotes the spread of polygenic host suppression.

To further unravel the differential selective pressure of the two CI outcomes on suppressor frequencies, let us first consider the diagram in Supplementary Figure S7A. Here, we pictorially describe the likely reason behind the differential increase in frequency of suppressors under CI. In a haplodiploid population with 50% frequency of suppressors, the expected number of suppressor alleles among the haploid oocytes of a female is 0.5*NL*, where *N* is the number of oocytes produced and *L* the number of loci (5 in our simulations). If an uninfected female oviposits 5 eggs, then the expected number of suppressors is 12.5. Upon fertilization by infected sperm carrying five active suppressors, CI is fully suppressed and the expected number of suppressors among offspring rises to *NL*[1.5*λ* + 0.5(1 − *λ*)], or 72.2% of all offspring loci for *λ* = 0.8 in the example, irrespective of the CI phenotype. Indeed, full suppression of CI does not distinguish between a MD-CI or a FM-CI regime, which are only detectable under the expression of the incompatibility.

The difference between MD-CI and FM-CI on the spread of suppressors lies in those CI crosses that involve males lacking suppressors. While MD-CI maintains the maternal alleles, including active suppressor alleles, in the population, all fertilized eggs die under a FM-CI regime, hereby increasing the rate at which active suppressor alleles are lost from the population (Figure S6A). We therefore argue that the essential factor that determines the speed at which suppressor alleles spread within a haplodiploid population is the net gain in the frequency of alleles derived from the CI crosses. Fixation of suppressors depends on their rate of increase through CI crosses involving males that carry suppressors and the rate at which they are lost due to CI crosses involving males lacking suppressors (Supplementary Figure S7B). To verify this argument, we measured the average frequency of suppressors derived from CI crosses among the offspring generated at each day of the simulation (Figure 5A) and the frequency of suppressors that are lost in crosses where CI is expressed (Figure 5B). Consistent with our expectations, we observed a continued marked difference between MD-CI and FM-CI. This confirmed that FM-CI causes a higher rate of information loss compared to MD-CI, and drives suppressors to fixation at lower rates. In Text S2 we further analyze the effect of fitness costs of suppressors to male hosts and show that, in all scenarios, infection persistency is always more stable under FM-CI.

**Figure 5.**
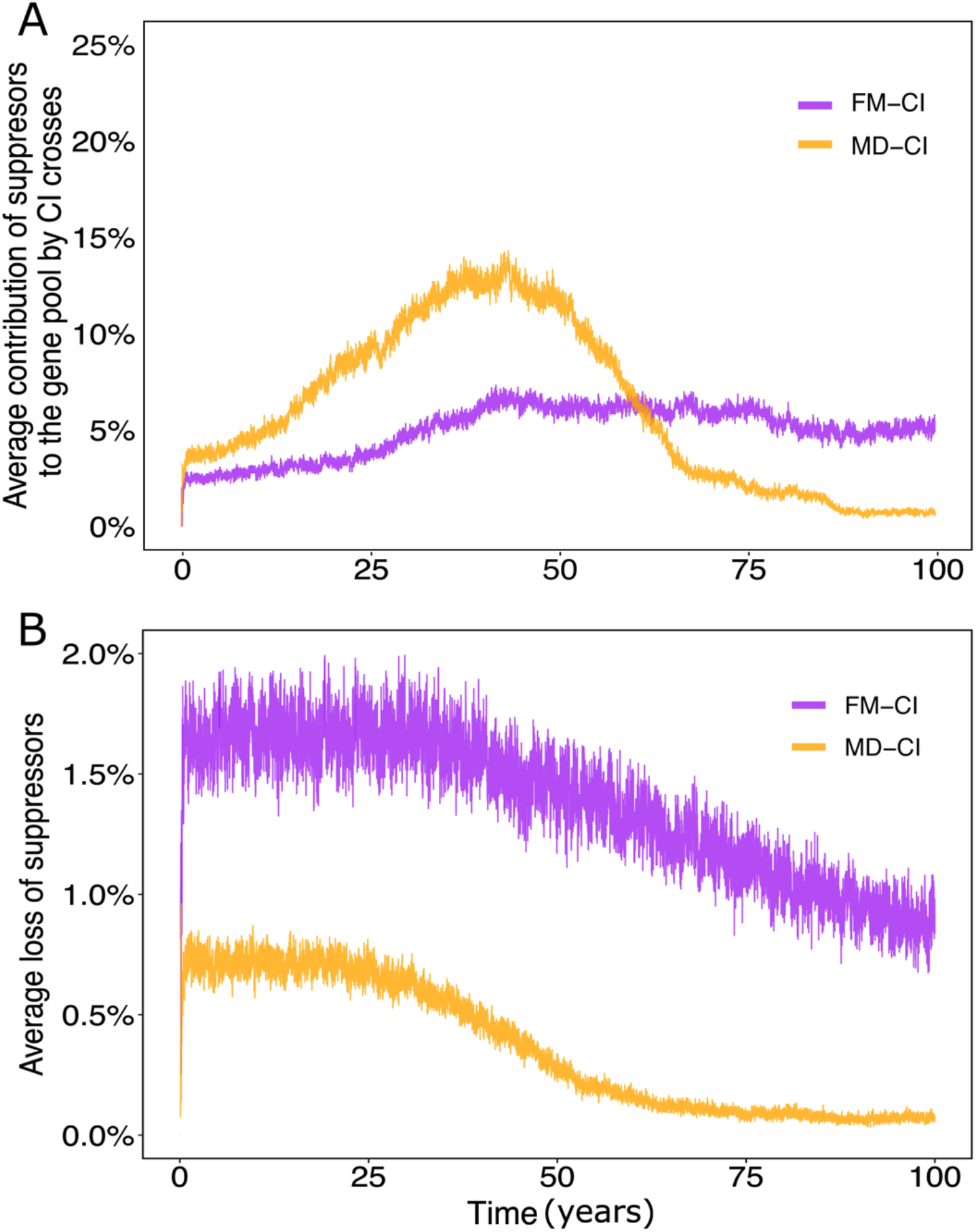
Effects of FM-CI and MD-CI on the spread and loss of host suppressors. Panel A depicts the average frequency of suppressors derived from incompatible CI crosses among all offspring generated at the end of a day. It is measured by counting the number of suppressor alleles in *new-borns* that derive from incompatible CI crosses divided by the total number of suppressors across all *new-borns*. Panel B represents the frequency of suppressors lost in an incompatible CI cross relative to compatible crosses, factored by CI type.

The long timescale for the spread of host suppressors indicates that the causal driving factor is the imperfect transmission rate *μ* of *Wolbachia*, as also suggested by previous theoretical work (Koehncke et al., 2009; Turelli, 1994). Imperfect transmission maintains a low frequency of CI crosses in the population despite that *Wolbachia* are at near-fixation. For a CI-inducing *Wolbachia* variant with perfect transmission (*μ* = 0) (CI parametrized according to *T. urticae* Beis), we observed that suppressor alleles remain centered at ∼50% and infection remains at fixation at all times (Figure 6A, B). This confirms the role of imperfect maternal transmission in the spread of polygenic host suppression. We further found that an imperfect transmission rate of 5% is sufficient to drive *Wolbachia* to extinction within the short span of ∼50 years in the presence of an initial frequency of 50% of suppressor alleles (Figure 6B). Finally, simulations with imperfect transmission rates ranging from 0% to 5% uncovered that the impact of this symbiotic trait on the spread of host suppressors differed significantly between FM-CI and MD-CI regimes (Figure 6C). Under a strict MD-CI regime, the spread of host suppression responded more sensitively to small imperfect transmission rates compared to FM-CI. This was interpreted as an expected result based on the higher efficiency of MD-CI as a driver of host suppression. This further supports the observation that reproductive symbiosis is more stable under FM-CI compared to MD-CI.

**Figure 6.**
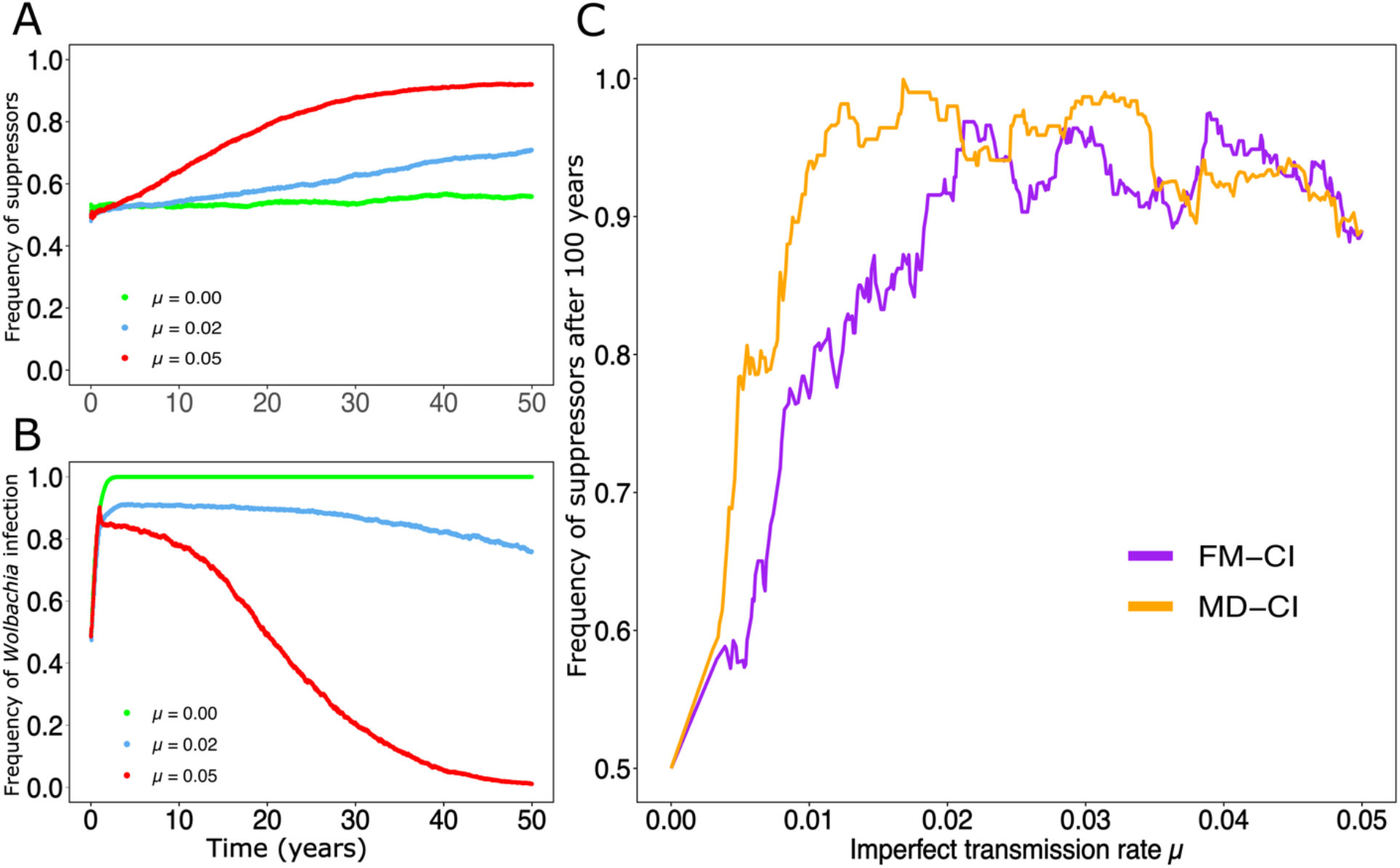
Effect of imperfect maternal transmission on the reciprocal spread of host suppression and *Wolbachia* infection. Panel A shows the effect of maternal transmission on the temporal changes in suppressor allele frequency. For *μ* = 0, we observe negligible changes, caused primarily by drift, in the frequency of suppressors. Increases in imperfect transmission rates led to a faster spread of host suppressors. CI was parametrized using *T. urticae* Beis. Panel B depicts the corresponding frequency of *Wolbachia* infection. Panel C shows the frequency of suppressor alleles measured after 100 years as a function of imperfect transmission rate. We distinguish between haplodiploid populations under FM-CI and MD-CI regimes.

### Sd extends *Wolbachia* persistence upon CI loss and facilitates mutualism

Sd is predicted to promote the spread of infection in simplified haplodiploid populations (non-overlapping generations and infinite population size) from low initial infection frequencies (Wybouw et al., 2023). To quantify the effect of Sd on *Wolbachia* spread in more realistic haplodiploid populations, we compared the infection profile of the population across the full range of *σ*_*total*_ and *ω*_0_ (with *σ*_*fm*_ = 0.98 and *σ*_*md*_ = 0.02). We found that Sd only exhibits an appreciable effect on *Wolbachia* spread when CI is weak (*σ*_*total*_ → 0). Final infection frequencies were increased by only a maximum of ∼8 and ∼10% for 3% and 5% Sd, respectively. The most pronounced effects of Sd were observed around intermediate values of starting infection frequencies (Figure 7A, B).

**Figure 7.**
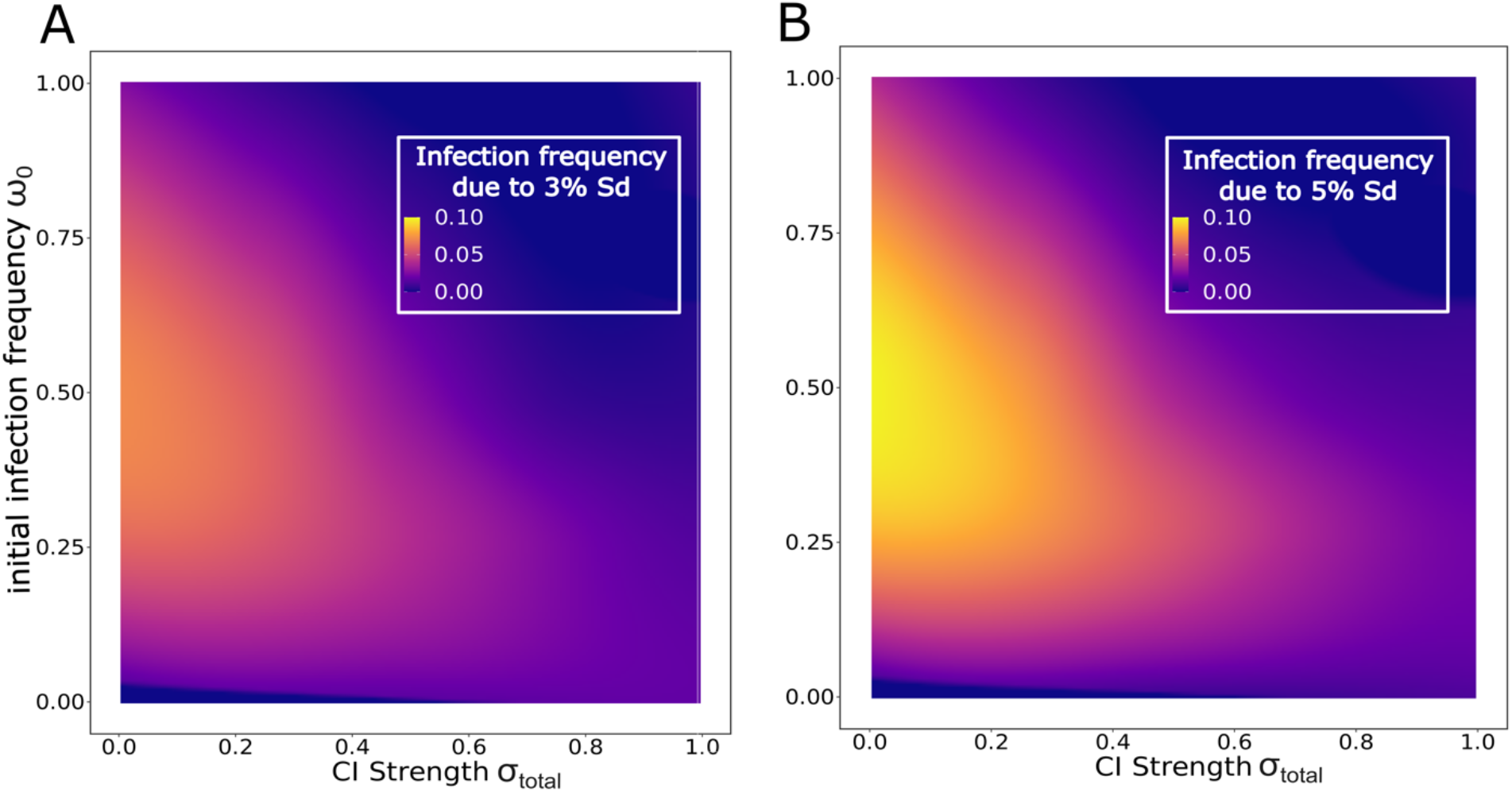
The effect of Sd on *Wolbachia* spread is most prominent with weak CI and intermediate initial infection frequencies. Both panels depict the increase in infection frequency by the end of a reproductive year when *Wolbachia* induce 3% and 5% Sd (panel A and B, respectively). Effect of Sd on final infection frequency was isolated by subtracting the infection profile of a population with Sd-inducing *Wolbachia* from the infection profile of a population without Sd. The relative proportions of MD-CI and FM-CI were experimentally derived from the Beis genotype (MD-CI 2%, FM-CI 98%).

In addition to CI loss by spread of host suppression, previous studies have shown that CI can be lost via pseudogenization of the *Wolbachia cif* repertoire (Meany et al., 2019; Prout, 1994; Turelli et al., 2022). These CI dysfunction events might be coupled with a transition towards a mutualistic infection through the evolution of positive female fecundity effects (Turelli et al., 2022; Weeks et al., 2007). Upon CI loss (*σ*_*total*_ = 0), we first quantified how long a neutral *Wolbachia* infection (no fecundity effects) persists in a haplodiploid population. We found that under CI loss, *Wolbachia* cannot maintain non-zero infection frequencies indefinitely when the system runs over many years. Moreover, the rates with which *Wolbachia* are lost depend on the magnitude of drift, imposed to the system at the transition of one year to the next (Supplementary Figure S8). In the absence of drift (i.e., when the entire population at the end of a reproductive year is carried over as the initial population of the next year), time to infection loss is well described by a gamma distribution, with an expected infection persistency of 15 years for a neutral *Wolbachia* infection (Figure 8A). When these simulations are replicated with *Wolbachia* variants that mediate 3% Sd, infection persistency is extended to 296 years (Figure 8A), a near 20-fold extension of infection persistency.

**Figure 8.**
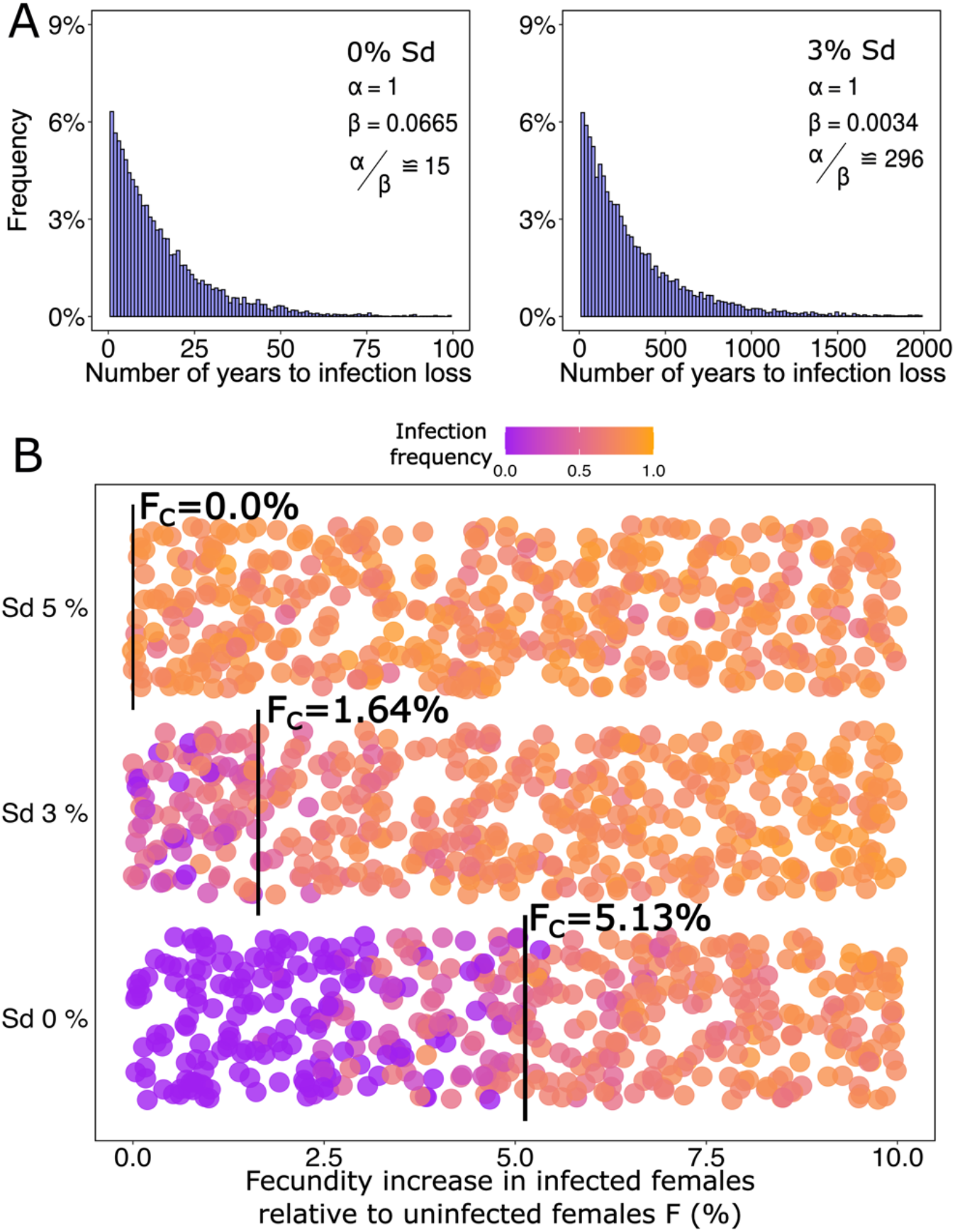
Upon loss of CI, Sd extends infection persistency and facilitates a mutualistic symbiosis. In panel A, we modelled infection persistency (number of years to infection loss) in the absence of CI (*σ*_*total*_ = 0) with 0% and 3% Sd. Infection persistency is well described by a Gamma distribution (parameters are outlined in the inset). Initial infection frequency was set as *ω*_0_ = 1. Panel B depicts the effect of relative fecundity increase (F) on infection frequency after 100 years factored by the strength of Sd. Vertical black bars indicate the critical F values. Here, *ω*_0_ = 0.5 and infection was considered stable or spreading when final infection frequencies were on average greater than or equal to *ω*_0_. In Figure S10, we further study the effect of Sd across a wider range of natural population sex ratios, *λ* ∈ [0.2, 0.8].

To study the transition into a mutualistic symbiosis, we quantified the fecundity threshold that allows *Wolbachia* to stably persist in the population upon CI loss. We measured the infection frequency after 100 years for various values of fecundity increase *F* ∈ [0, 0.10], sampled from a uniform distribution, using ω_0_ = 0.50 and *σ*_*total*_ = 0 (where the effect of Sd on final infection frequency is the strongest, Figure 7). We ordered the obtained infection frequencies with respect to a non-decreasing sequence of *F* and computed the moving average of infection frequency using a lag period of 10 observations. The threshold value of fecundity (*F*_*c*_) was taken to be the value of *F* for which the average infection frequency was greater than or equal to ω_0_ (Supplementary Figure S9). For a *Wolbachia* variant that does not induce Sd, stable infections can only be attained with a fecundity increase of ∼5.13%. Replicating these simulations with Sd-inducing *Wolbachia* variants uncovered that Sd considerably reduces *F*_*c*_ (Figure 8B). For Sd = 3%, we identified an approximate three-fold decrease in fecundity necessary for stable infection persistence (∼1.64%). For Sd = 5%, there was no fecundity benefit required to maintain a stable *Wolbachia* infection. A lower-bound for Sd strength necessary to maintain infection stably in the system can be obtained by Equations 6 − 7. For ω_0_ = 0.5 and F = 0% Equation 6 must satisfy:

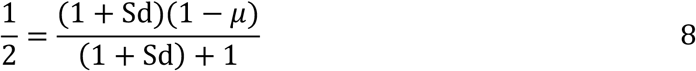

which for *μ* ≈ 0.02, according to *T. urticae* data, yields,

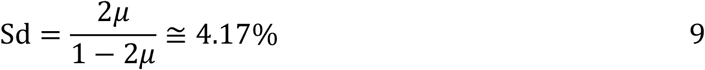

More generally, the minimum Sd strength necessary to maintain any non-zero initial infection frequency ω_0_ stable can be computed by considering the following relation derived from Equation 6 with *F* = 0:

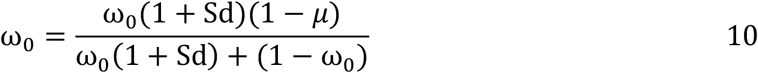

Assuming ω_0_ > 0, Sd reads:

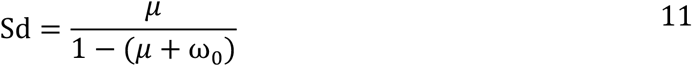

Because Sd is a non-negative quantity, Equation 11 is defined only when (*μ* + ω_0_) < 1. Thus, within the open interval (0, 1), it describes a family of curves parametrized by *μ* which ultimately determines the strength of Sd necessary to stably maintain ω_0_ (Figure 9A). Moreover, if Sd is non-zero, we arrive at:

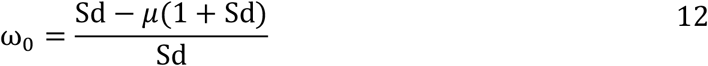

Thus, upon CI loss, one may predict the infection frequency that is stably maintained in the system due to Sd and *μ* alone using Equation 12. We confirmed these stable polymorphic infection frequencies with our individual-based simulations (Figure 9B). Note that while Equation 12 does not depend explicitly on λ, Sd does, with its maximum magnitude given by 1 − λ.

**Figure 9.**
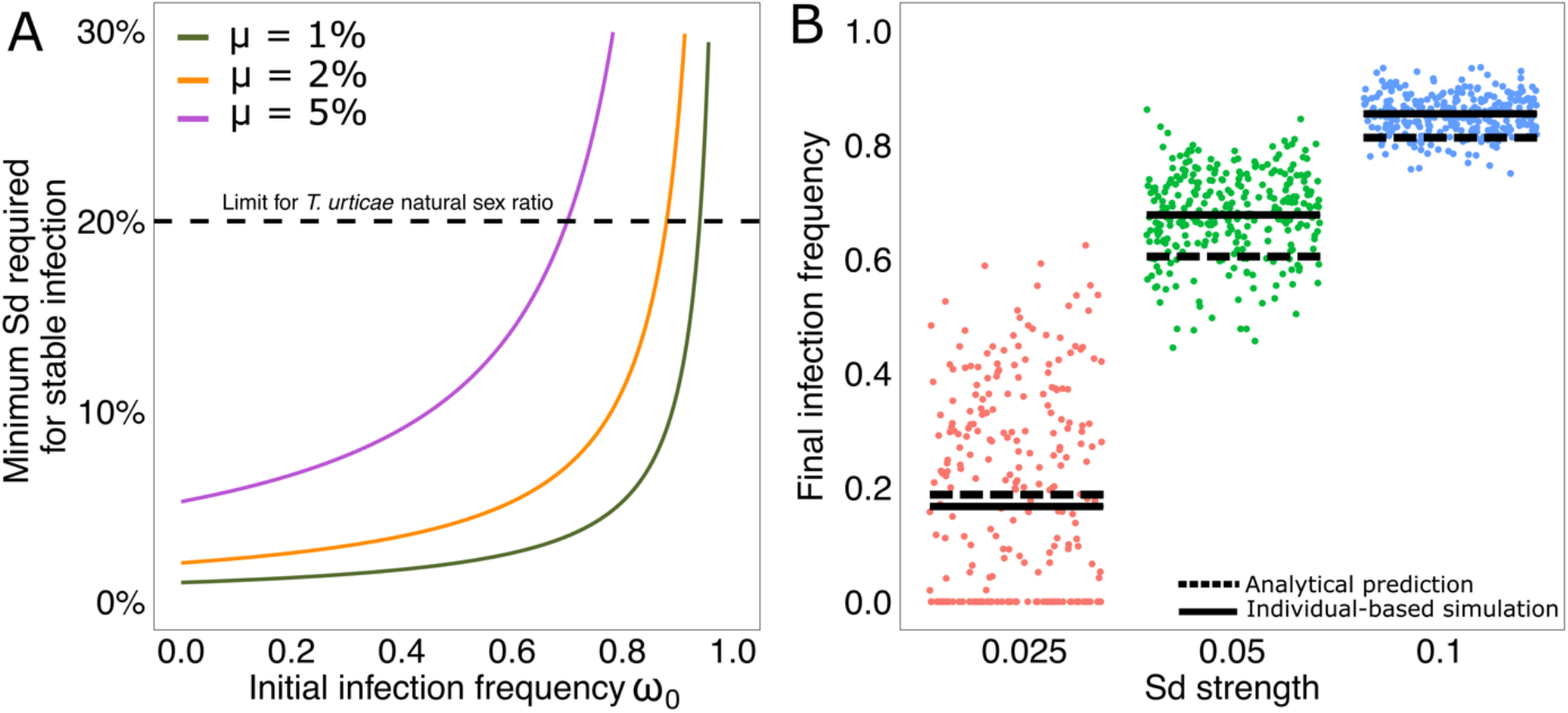
Weak Sd can maintain a stable polymorphic infection frequency upon CI loss. In panel A, we show the relationship between the strength of Sd and maternal transmission to keep a constant infection frequency for any given initial infection frequency *ω*_0_. Three curves are displayed for different imperfect transmission rates *μ*. A horizontal dashed line depicts the average maximum strength of Sd for Beis *T. urticae*. In panel B, the infection frequencies after 10,000 days of the populations in the individual-based simulations are depicted. For each value of Sd in the x-axis we plot the results of the IBM (average as solid black lines and individual simulations’ observations as dots) and the predicted infection frequencies by Equation 12.

## Discussion

The widespread occurrence of reproductive symbionts in nature and the relevance of *Wolbachia-* based pest control necessitates a thorough understanding of the mechanisms that underlie infection persistency, including in hosts that have a haplodiploid mode of reproduction. Our simulations demonstrated that *Wolbachia* that induce complete CI in haplodiploids only require threshold infection frequencies of ∼20% to reach near-fixation within ∼15 host generations, regardless of CI phenotype (MD-CI or FM-CI). Threshold infection frequencies for the spread of *Wolbachia* are central to *Wolbachia-*based pest control and have commonly been associated with fitness penalties of infection such as reduced female fecundity (Hoffmann et al., 2011; Schmidt et al., 2017; Turelli et al., 2022; Turelli & Barton, 2017, 2022). In contrast to previous theoretical study (Turelli, 1994; Turelli et al., 2022; Vavre et al., 2000; Wybouw et al., 2023), threshold infection frequencies manifested in our simulations without implementing fecundity costs of infection, but as a result of variance that emerges from the dynamics of populations with finite size and overlapping generations. These findings caution to explain observed bistable *Wolbachia* dynamics in field populations by fecundity effects alone. Moreover, this observation further underscores that estimated fecundity effects based on experimentation with laboratory populations cannot fully predict natural *Wolbachia* dynamics in field populations.

The developmental outcome of CI in haplodiploids lies on a phenotypic continuum with complete FM-CI and MD-CI as the extremes. Our findings are consistent with previous theoretical work and predict that MD-CI should be less efficient in driving *Wolbachia* infection compared to FM-CI (Vavre et al., 2001). Our high-resolution quantification of the driving efficiencies verified that, within a year, MD-CI can be up to ∼12% less efficient than FM-CI. In agreement with the arguments developed by previous work assuming monogenic CI suppression and a constant infection frequency (Vavre et al., 2003), we further show that host suppressors spread ∼2.5-fold faster under MD-CI compared to FM-CI. The spread of polygenic suppression coincided with the rapid loss of CI-inducing *Wolbachia* in either case. These divergent host-symbiont dynamics in response to CI type are explained here by different rates of genetic information loss in infected haplodiploid populations. The retention of suppressor alleles in the population accelerates the spread of host suppression under an MD-CI regime. Together, our findings that MD-CI dramatically reduces infection persistency by decreasing *Wolbachia* drive efficiency and promoting the spread of suppressors raise the question of why both CI types are co-expressed and maintained in haplodiploid populations/species (Mouton et al., 2005; Perrot-Minnot et al., 2002; Vavre et al., 2001; Wybouw et al., 2022). Our ability to address this question is still limited because the genetic factors that determine whether a CI embryo dies (FM-CI) or develops as a male (MD-CI) are not fully resolved. Studies on *Nasonia* and *Tetranychus* show that the female genotype modulates CI type and the maternal genetic factor that causes strong MD-CI is recessive (Bordenstein et al., 2003; Wybouw et al., 2022). The developmental outcomes of CI crosses are also modulated by the symbiotic species. In a coinfected *Pezothrips* population, MD-CI was linked to *Cardinium* but not to *Wolbachia* (Nguyen et al., 2017). Future study should formally test whether *cif* gene repertoires might shape the developmental outcome of *Wolbachia-*induced CI in haplodiploids.

In natural populations CI is often weak or not detectable (Shropshire et al., 2022; Wybouw et al., 2022), observations that are typically coupled with the development of host suppression and/or a mutualistic association between *Wolbachia* and its host (Egas et al., 2002; Prout, 1994; Weeks et al., 2007). Based on our current mechanistic understanding of host effects on CI strength, we assumed an additive polygenic basis for host suppression of CI in *T. urticae* (Cooper et al., 2017; Reynolds & Hoffmann, 2002; Wybouw et al., 2022). Under this assumption, we demonstrate that the selective pressure of CI during a *Wolbachia* invasion event does not lead to an appreciable increase in suppressor allele frequency. Instead, the residual low CI cross frequency, caused by imperfect maternal transmission, exerts the positive selective pressure on the host for the spread of polygenic host suppression on longer timescales (>500 host generations). We observe that empirically derived transmission rates are sufficient for the spread of suppressors in *T. urticae* populations. We found that imperfect transmission rates as low as 0.5% (well below our experimentally derived estimate of ∼2%) are sufficient to promote the spread of suppressors and the loss of infection. Moreover, MD-CI was more sensitive to imperfect transmission compared to FM-CI, which follows from their increased efficiency to promote the spread of host suppressors. As the maternal transmission rates of *Wolbachia* negatively covary with temperature in *Tetranychus* mites (Van Opijnen & Breeuwer, 1999), we expect temporal effects on the spread of suppression in natural populations. In addition to transovarial maternal transmission, *Wolbachia* are also horizontally acquired, albeit at low frequencies (Sanaei et al., 2021; Turelli et al., 2018). For arthropod herbivores such as *Tetranychus* mites, host plants may act as a transmission route and reservoir (Li et al., 2017; Sanaei et al., 2021). Whether rare horizontal transmission can modulate infection persistency and the spread of suppression is not understood and is an interesting avenue for future research.

To quantify infection persistency upon host suppression, we studied unidirectional CI (i.e. uninfected females crossed to infected males) by introducing a single CI-inducing *Wolbachia* variant in our populations. However, CI can also be bidirectional when mutually incompatible *Wolbachia* variants co-segregate in a host population. For some systems, results indicate that bidirectional CI occurs when CifA proteins do not bind and neutralize CifB proteins from the other variant (Namias et al., 2022; Wang et al., 2022; Xiao et al., 2021). In *Drosophila teissieri*, an interaction between host and *Wolbachia* genetic variation determines CI strength (Cooper et al., 2017), suggesting that host suppression might act differently on divergent Cif proteins. It is tempting to speculate that polygenic host suppression might modulate bidirectional CI in complex ways, creating highly unpredictable infection persistency in natural populations.

In our model, neutral *Wolbachia* (no effect on fecundity and upon loss of CI either via host suppression or *cif* pseudogenization) were consistently purged from haplodiploid populations within ∼20 years, a loss that was (mainly) driven by *Wolbachia* imperfect transmission. Upon CI loss, we showed that a relative fecundity increase in infected females of ∼5% is sufficient for a stable symbiosis. This threshold fecundity increase was surprisingly low since experimental work on *T. urticae* field populations uncovered relative fecundity benefits of up to ∼25% (Zélé et al., 2020). Transition to mutualism in field populations has been documented in *Drosophila simulans*, a diplodiploid insect, by Weeks et al. (2007). In the focal *D. simulans* populations, *Wolbachia* promote a relative fecundity increase of ∼10% within the span of 20 years of invasion. However, in our study, only the effect of infection on female fecundity was modelled, ignoring potential fitness trade-offs. *Wolbachia* are known to modulate other fitness components, including longevity, in *Tetranychus* mites and trade-offs between fecundity and longevity have been observed (Vala et al., 2002; Zélé et al., 2020). For *Wolbachia* variants that induce 3% Sd, we found that infection remains in a haplodiploid population for ∼300 years without relative fecundity benefits, approximately a 20-fold increase in infection persistency relative to the absence of Sd. This remarkable difference in infection persistency reduces the fecundity threshold to ∼1.6% for a stable mutualistic association. These results broadly confirm our hypothesis that Sd may be an important component in driving *Wolbachia* symbioses from a parasitic to a mutualistic association in haplodiploids. Here, to study the development of stable mutualism, we assumed a complete simultaneous loss of both CI induction and rescue by pseudogenization of entire *cif* operons. Studies have however shown that the ability of *Wolbachia* to rescue CI can be maintained after loss of CI induction (Bourtzis et al., 1998; Vala et al., 2002). This is currently explained by higher frequencies of pseudogenized *cifB* genes, the single causal genetic factor of CI induction in most systems (J. F. Beckmann et al., 2021; Martinez et al., 2021) (but see (Shropshire & Bordenstein, 2019)). Theory predicts that rescuing variants that do not induce CI can still spread if CI-inducing variants infect the population at high frequencies (Hurst & McVean, 1996; Prout, 1994; Turelli, 1994). As arthropod species are often infected by multiple *Wolbachia* variants (Kriesner et al., 2013; Wolfe et al., 2021), variation in the *cif* gene repertoire across co-existing *Wolbachia* variants could shape the evolution of mutualism.

For *Wolbachia* variants that induce 5% Sd, infection was stable and polymorphic without any relative fecundity benefits. Together with previous work (Egas et al., 2002; Wybouw et al., 2023), there is growing theoretical support that Sd alone can drive *Wolbachia* to near-fixation and maintain the infection at high frequencies. Experimental work that uncovers Sd levels of up to ∼18% further strengthen this conclusion (Vala et al., 2003). Although both host and *Wolbachia* are known to modulate the penetrance of Sd (Vala et al., 2003; Wybouw et al., 2023; Zélé et al., 2020), the causal genic factors remain unresolved.

