## Supplementary Material for "Reciprocal host-*Wolbachia* interactions shape infection persistence upon loss of cytoplasmic incompatibility in haplodiploids"

### Supplementary Material for "Impact of reproductive phenotypes on the reciprocal dynamics of *Wolbachia* and haplodiploid hosts"

### Text S1. Additional technical information on the implementation of simulations

Here, we describe the technical details concerning the implementation of the individual-based model (IBM) to study infection persistency of *Wolbachia* in haplodiploid hosts. The original implementation was carried out in Java SE 20 and is available in a public GitHub repository (*link-after-acceptance*). The aim of Text S1 is to provide further information on the general logical structure of the model to ease implementation in other programming languages. We highlight that the original code is available with an extensive number of comments where the Java-specific syntax may hinder the understanding of the reader. Naturally, there are a number of different choices when it comes to implementation of specific *Methods* and therefore readers may find the original code useful for this purpose.

We consider an IBM to simulate a haplodiploid population of initial size $P_0$ with overlapping generations. Individuals have an explicitly determined sex, age (measured in days), and *Wolbachia* infection status (infected or uninfected). Females are diploid and males are haploid. Individuals reach maturity at age $M$, after which females are considered adult and start to oviposit haploid eggs, whereas males are allowed to fertilize adult females. Females only need to be fertilized once, and will carry the sperm throughout their entire lives. Both females and males can live at most $D$ days. The program consists of updating an Initial population by iteratively erasing Individuals that reach age $D$ and inserting offspring that emerges from eggs oviposited by adult females, according to the rules discussed in the main manuscript. Egg hatching is modulated by the strength of FM-CI. If a fertilized egg suffers from MD-CI, the embryo will develop into a male, instead of a diploid female. Below we provide the pseudo-code (**Algorithm 1**) for the general logic of the model. *Methods* are highlighted in bold red italic, control statements are in bold black, parameter names are in italic and comments are given between forward slash and asterisks.

In the original code, female and male individuals are Objects with a certain number of attributes, such as current age, maturity age, maturity peak (for females only), age of death (defined upon egg hatching) and infection status (Boolean). Likewise, we conceptualize an egg as an Object which inherits the chromosome (linear vector) of both parents if fertilized, or only the maternal chromosome if no fertilization occurs. The *Egg* contains methods that control whether it hatches or not, as well as the probability that it is infected by *Wolbachia* when it is created by an infected female. These Objects are stored in lists that represent the entire population, where operations such as calculation of infection and suppressor frequencies are performed. Population level methods are described in full detail in the original code, where they are stored inside a *Class* called *PopulationLevel*.

When one wishes to simulate the dynamics of infection across multiple years, **Algorithm 1** must be wrapped inside a *while* loop that controls the number of years. After each execution of **Algorithm 1** the system must choose a fraction of individuals inside *initPop* that will make up the initial population of the next year. We perform this operation by retrieving adult individuals (both males and females) from *initPop* randomly, i.e., each adult has equal chances of becoming part of the founding population of the next year. One must also consider that Equation 2 from the main manuscript must be adapted accordingly. That is, for a given carrying capacity K, the initial population size in the equation must be changed flexibly to avoid that *Drift()* does not operate on the population following the expected population size as determined by $P_0$.

**Algorithm 1. Host – symbiont dynamics during a single year.**

```
initPop ← P_0 /* Initialize population */
days ← 150 /* Number of days in a growing season */
t ← 1 /* Counter to control time */
while t ≤ days do
    newPop ← 0 /* Initialize empty population that will store offspring for the next day */
    adultFemales ← findFemales(initPop) /* Find all female individuals in reproductive age from initPop */
    adultMales ← findMales(initPop) /* Find all male individuals in reproductive age from initPop */
    for i in adultFemales do
        numEggs ← 0 /* Initialize number of eggs laid by female i */
        fertilized ← checkFertilization() /* Check whether female i is fertilized */
        if fertilized then
            numEggs ← oviposit() /* Number of oviposited eggs (see below and Equation 1 in the main manuscript) */
        else
            male ← random(adultMales) /* Choose random male to fertilize female i if adultMales is not empty */
            fertilize(adultFemales[i], male) /* Fertilize female i with sperm from male */
            numEggs ← oviposit()
        end if
        for j in numEggs do
            egg ← hatching() /* Compute probability egg becomes diploid and hatches (see below) */
            egg ← infectionStatus() /* Compute the probability that Wolbachia is transmitted to the egg */
            if egg is diploid and hatches then
                newPop ← newPop + egg /* Egg contains maternal and paternal genetic material */
            else if egg is haploid and hatches then
                newPop ← newPop + egg /* Egg contains only maternal genetic material */
            end if
        end for
    end for
    increaseAge(initPop) /* Every individual gets older by one day and dies if age is equal to or greater than D */
    initPop ← initPop + newPop /* Initial population receives original aged population and newly formed individuals */
    drift(initPop) /* Apply drift to keep population under expected size (see Equation 2 in the main manuscript) */
    t ← t + 1 /* Go to next day */
end while
```

The method *oviposit()* in **Algorithm 1** determines the number of eggs laid by a female based on its natural fecundity and *Wolbachia*-induced increase in fecundity (if F>0). Then, the number of eggs to be fertilized is determined according to $\lambda$ and *Sd*. The natural fecundity of a female is a function only of its current age (see Equation 1 in the main manuscript). Oviposition operates as in **Algorithm 2**.

Note that for each egg laid, *F* determines the probability that an additional egg is generated. The expected number of additional eggs is the number of eggs of the original batch multiplied by the probability of a new egg generated per egg oviposited. So, if $F = 0.10$, there will be 10% more eggs than the original batch. Sex allocation distortion operates in the same manner, but increases the expected number of eggs that receive sperm, and are thus diploid. In the original Java implementation, the number of diploid eggs is determined before the *for* loop in which *hatching()* operates. Then, *hatching()* only calculates the probability that a diploid egg will hatch according to the rules of the main manuscript or turn into a haploid egg if $\sigma_{md} > 0$.

**Algorithm 2. Oviposit**$(adultFemales[i], age, F, Sd)$

```
eggs.size() ← getFecundity(age) /* get number of eggs that a female with current age "age" should oviposit */
eggs.genotype() ← receives only haploid maternal chromosome /*Random loci from female's genotype */
for i in eggs.size() do
    r ← random() /* Generate random number with uniform distribution between 0 and 1 */
    if r < F then
        eggs.size() ← eggs.size() + 1
    end if
end for
/* determine eggs that shall be fertilized */
for i in eggs.size() do
    r ← random() /* Generate random number with uniform distribution between 0 and 1 */
    if r < λ then
        eggs.genotype(i) ← adultFemales[i].sperm /* egg becomes fertilized, i.e., diploid */
    end if
end for
for i in eggs.size() do
    if eggs.genotypes(i) is haploid then
        r ← random() /* Generate random number with uniform distribution between 0 and 1 */
        α ← Sd/(1 − λ) /* According to Equation 5 in the main manuscript */
        if r < α then
            eggs.genotype(i) ← adultFemales[i].sperm /* egg becomes fertilized, i.e., diploid */
        end if
    end if
end for
return eggs
```

### Text S2. Analysis of impact of fitness costs on the spread of polygenic host suppression

Whether CI suppressors correlate with a fitness cost in male hosts is unknown, but theory suggests that, under certain conditions, monogenic suppression that causes fitness penalties could still spread within host populations and lead to a loss of *Wolbachia* infection (Koehncke et al., 2009). Previous experimental work in *Drosophila melanogaster* found that resistance to the intestinal pathogen *Pseudomonas entomophila* correlates with lower sexual competitiveness among male hosts in susceptible populations (Kawecki, 2020). Such studies often generate contrasting results on the relationship between sexual competitiveness and microbial resistance (McKean & Nunney, 2001; McNamara et al., 2013; Rolff & Kraaijeveld, 2003; Ye et al., 2009). Here, we further tested the effect of fitness penalties of active suppressors to male hosts. Specifically, we investigated whether a fitness cost to male sexual competitiveness had any appreciable effect on the spread of suppressors in haplodiploid populations. The issue arises when one asks whether sexual fitness costs have an appreciable effect on the spread of suppressors depending on CI phenotype, i.e., MD-CI versus FM-CI. Therefore, we introduced a parameter $\phi \in [0, 0.02]$ which represents the fitness cost of each active suppressor in the male haplotype. If $L_s$ is the number of active suppressors in a randomly chosen male for reproduction, then $\phi L_s$ corresponds to the probability that another male is picked among the males in reproductive age to fertilize a certain virgin female. The probability measure $\phi L_s$ is well defined as $\max[L_s] = 5$ (see main text) and $\max[\phi] < 0.2$.

In the figure below, we depict the effect of a reproductive cost on the spread of polygenic host suppression for MD-CI and FM-CI. For $\phi = 0.015$ , we found that suppressors were purged faster under FM-CI compared to an MD-CI regime (Panel A). Interestingly, for $\phi = 0.010$, we observed a spread of host suppression under MD-CI, whereas host suppression was still purged within the FM-CI regime. For $\phi = 0.005$, suppressor frequency did not rise to fixation within 100 years under an FM-CI regime, whereas an MD-CI regime quickly led to the fixation of active suppressor alleles. In general, for any reproductive cost associated with host suppression, suppressors spread more efficiently under MD-CI, destabilizing *Wolbachia* symbiosis (Panel B).

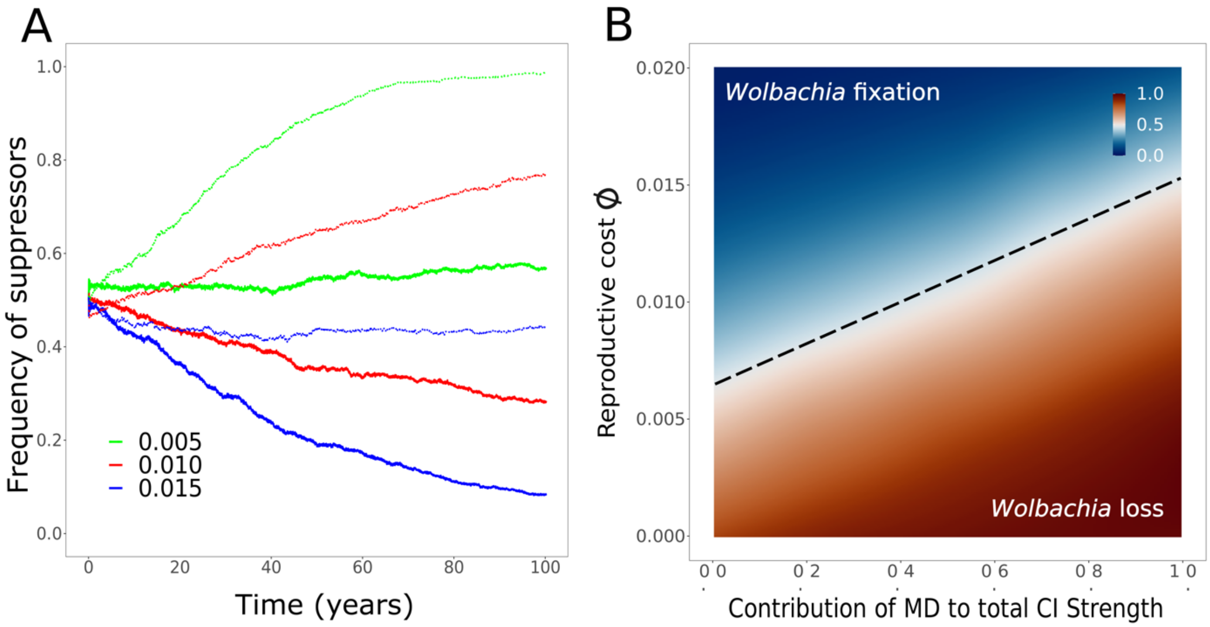

**The impact of fitness penalties on the spread of polygenic host suppression is contingent on the CI phenotype.** Panel A shows the frequency of suppressors in function of time (year) for different fitness costs $\phi$ (values are color coded and outlined in legend). Solid thick lines represent simulations with FM-CI whereas thin dotted lines represent an MD-CI regime. Panel B depicts the frequency of suppressors after 100 years as a function of the reproductive cost $\phi$ and the contribution of MD-CI to total CI strength ($\sigma_{fm} = 1 - \sigma_{md}$).

### Supporting Figures

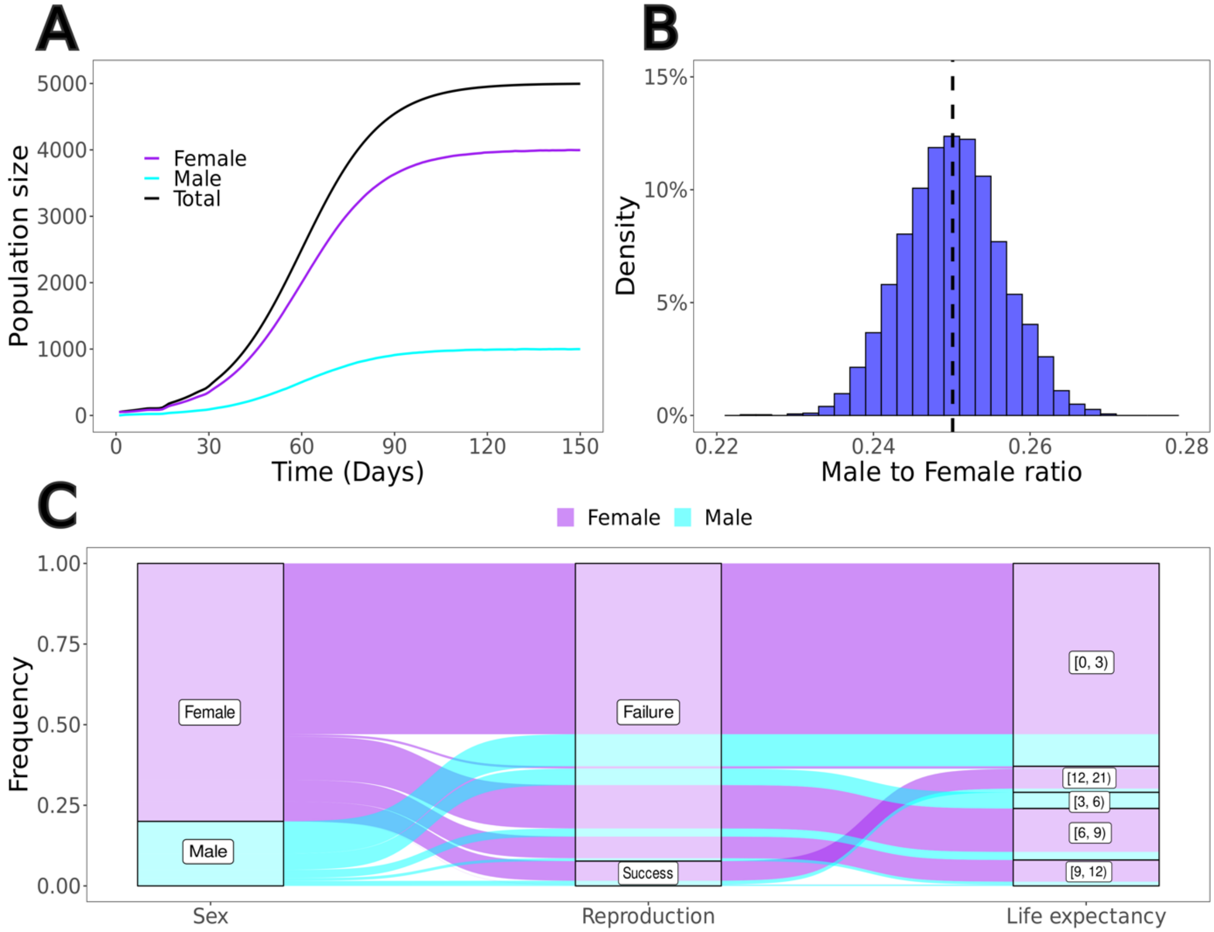

**Figure S1. General description of the demographic state of the simulation under a *Wolbachia*-free haplodiploid population.** Panel A depicts the population growth during a single year. The population reaches carrying capacity K=5000 by the end of the year with a normally distributed male to female sex ratio according to the parametrization of the model of $\lambda = 0.8$ as shown in panel B. In panel C, we show an alluvial plot with the general dynamics of the model. 6.2% of the females and 1.6% of the males reach adulthood and successfully reproduce. Life expectancy of reproducing individuals often exceeds 12 days. The vast majority of individuals die within 3 days. This is attributed to the drift component (Equation 2) that targets random individuals of the population irrespective of age and sex. Because at the end of each day, the number of eggs is larger than the number of adults, drift targets a higher proportion of newborns.

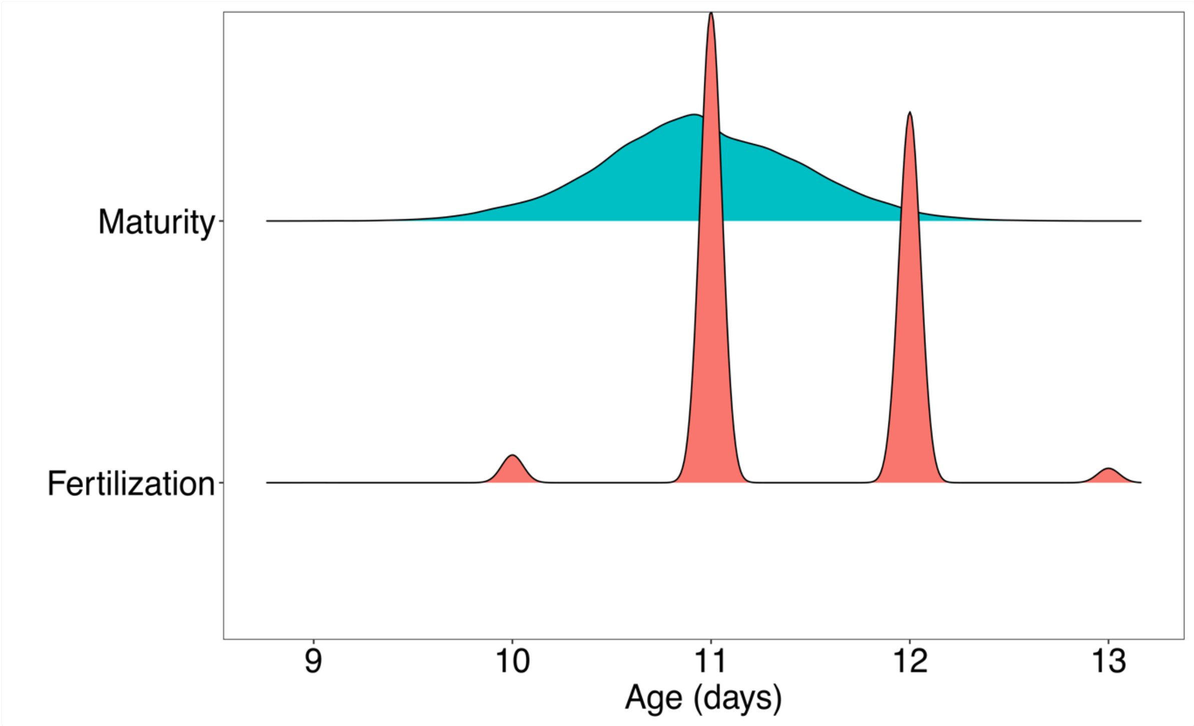

**Figure S2. Distribution of maturity and fertilization age of females.** The figure depicts the frequency distributions of the female maturity age (a parameter in the system) and the age at which virgin females are fertilized (an emergent property). Note that fertilization occurs approximately as soon as females reach maturity. Smooth distributions are shown for aesthetical purposes. Fertilization can only occur in discrete time units. That is, the smallest unit of time measured in our simulations is one day.

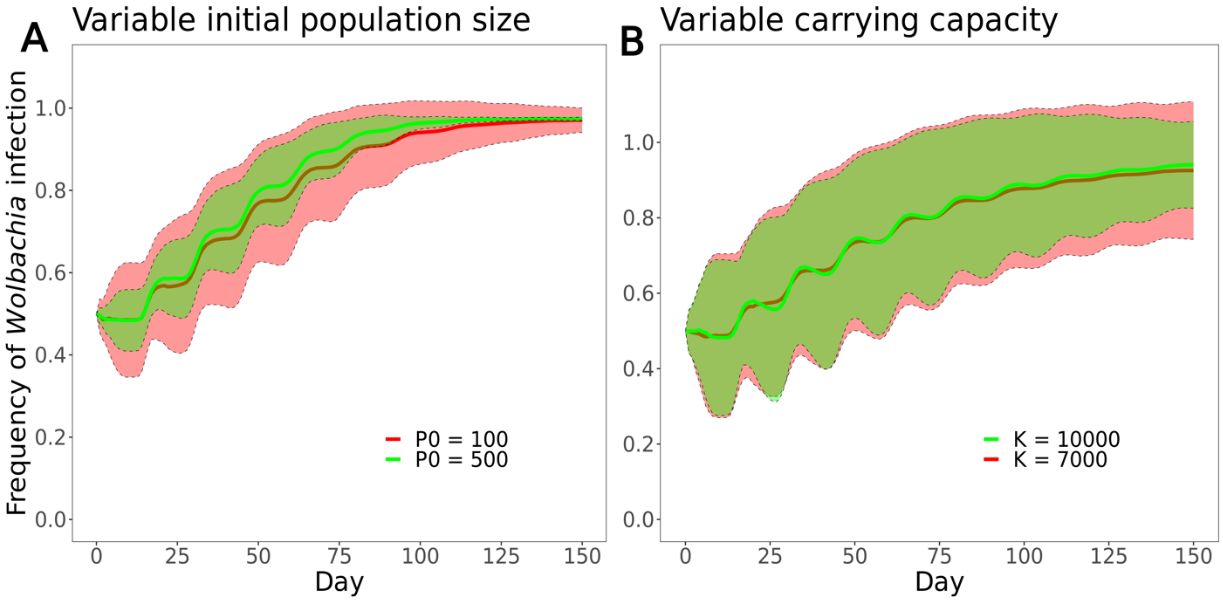

**Figure S3. Effects of initial host population sizes ($P_0$) and carrying capacities ($K$) on the spread of infection during a reproductive year.** For any starting and final population size, we verified that our IBM returned consistent predictions compared with the chosen parameters of $P_0$ and $K$ used in the main manuscript. Solid lines represent the average frequency of *Wolbachia* infection per day. Ribbons represent $\pm$ one standard deviation around the average. Note that for higher starting population sizes, the standard deviation decreases accordingly. The low $P_0$ used in our experimental setting might explain the high variance found in the outcome of the IBM within the main manuscript, which closely approximated the standard deviation of the experimental results. This effect is most likely a consequence of drift, which is expected to disappear in the limit of infinite population sizes.

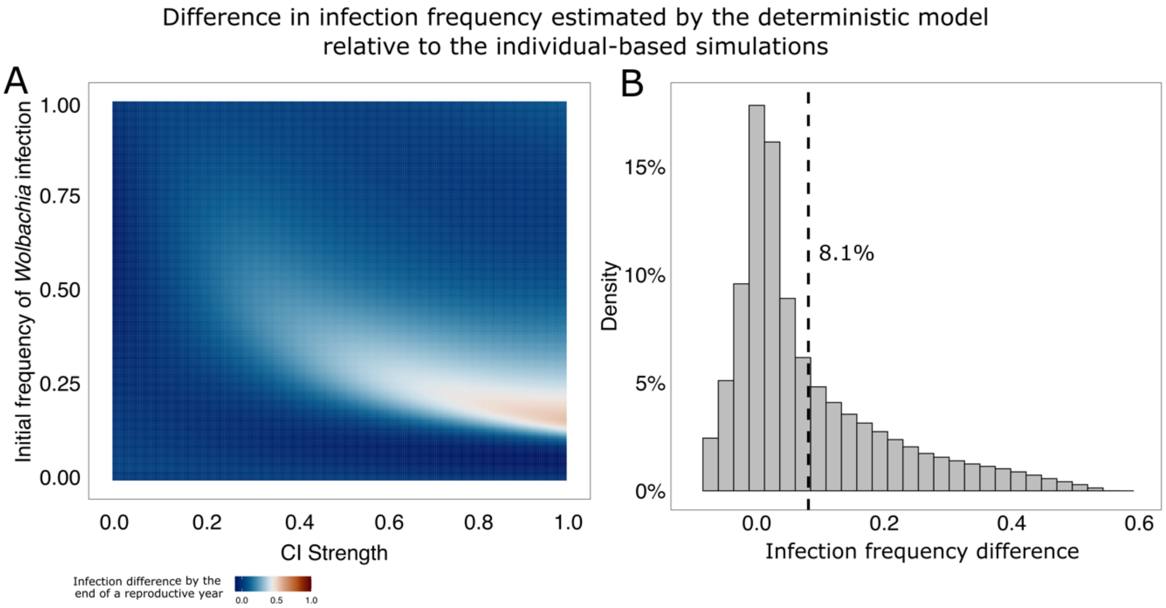

**Figure S4. Comparison between the deterministic model of infection spread in haplodiploids and the individual-based simulations built for *Tetranychus urticae*.** Panel A) depicts the difference in infection frequency by the end of a reproductive year (150 days) estimated by the deterministic recursions given by Equations 6-7 in the main manuscript relative to the estimates by the IBM. That is, infection frequency estimated by the recursion minus infection frequency estimated by the IBM, across the span of CI strength and initial *Wolbachia* frequencies. In Panel B) the frequency distribution of these differences is presented. The average difference was found to be 8.1%, which implies that the deterministic model overestimates infection frequencies in the population by 8.1% relative to individual-based simulations.

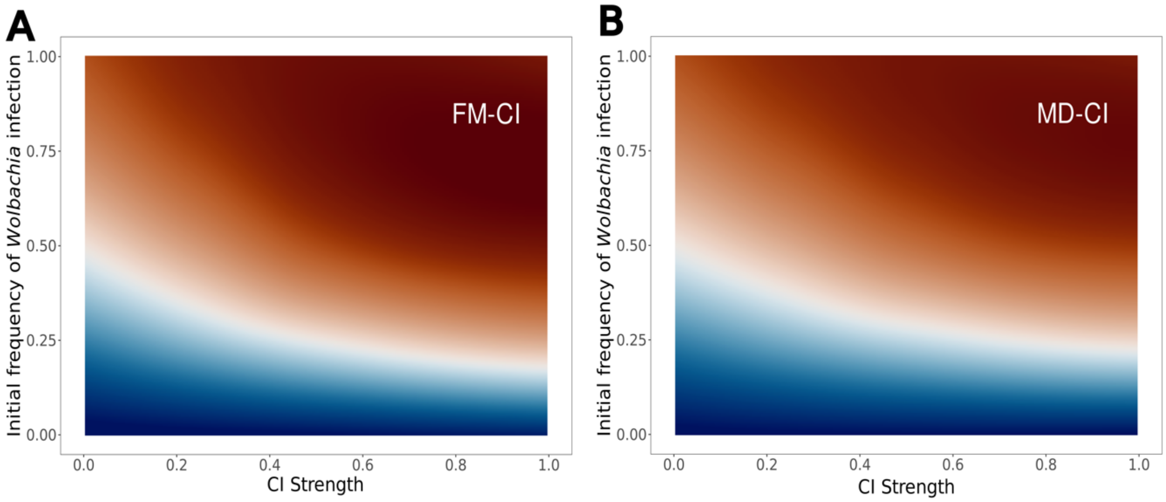

**Figure S5. Infection profile of haplodiploid populations under different CI regimes.** Panels A and B show the infection frequency by the end of a reproductive year for a population under FM-CI and MD-CI, respectively. The general pattern of infection frequencies across the full range of initial infection frequency ($\omega_0$) and CI strength ($\sigma_{total}$) shows no differences between the two CI types. However, we identified important quantitative differences that are discussed within the main manuscript.

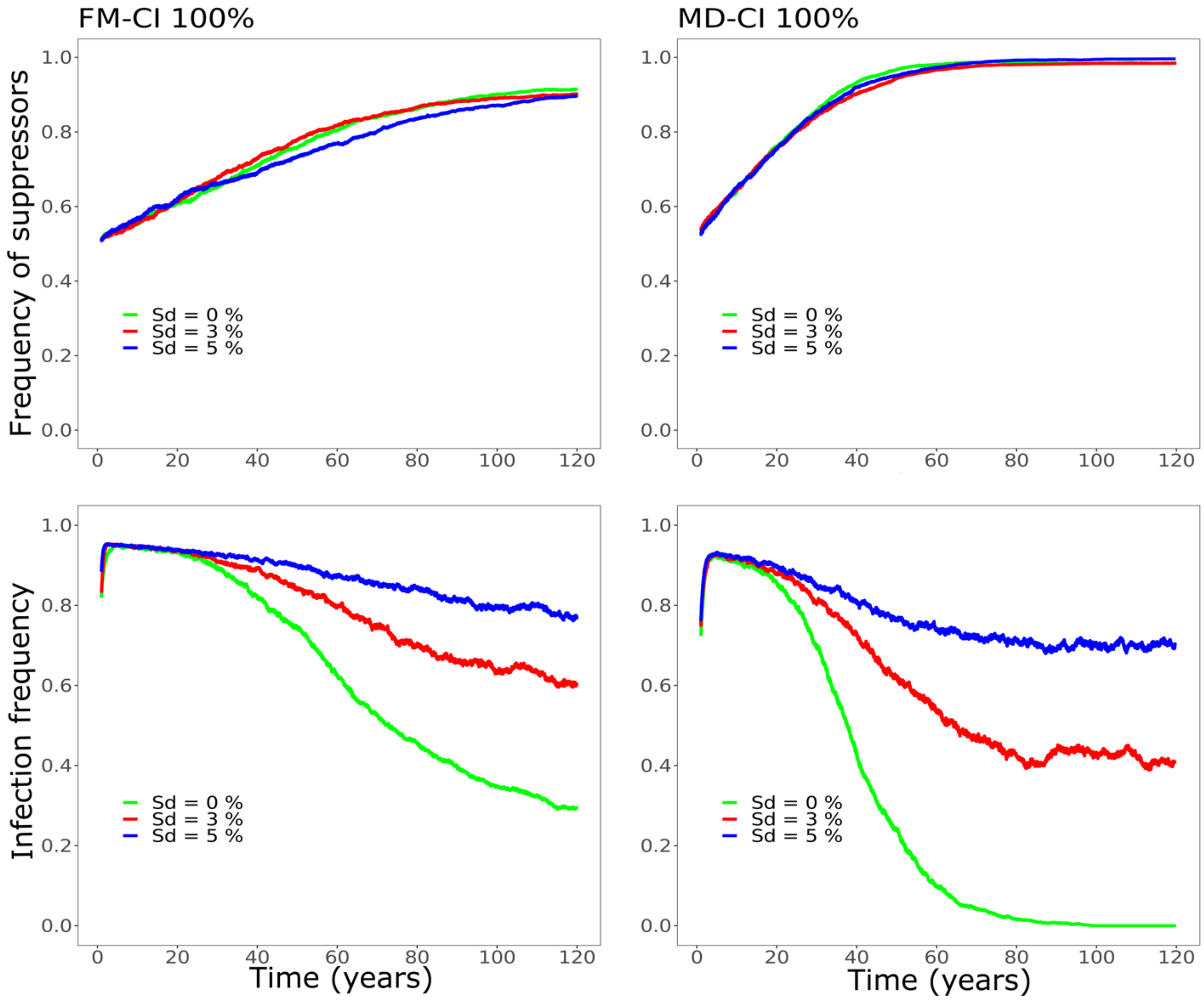

**Figure S6. Sex allocation distortion does not display any appreciable effect on the spread of CI suppressors.** Left-hand side panels show the frequency of suppressors and corresponding *Wolbachia* infection frequencies in a haplodiploid population for varying Sd strength in a FM-CI regime. Right-hand side panels show the same information for an MD-CI regime. Note that in neither case the spread of suppressors is affected by Sd. The effect of Sd on *Wolbachia* infection frequencies can only be observed after suppressors have reached near-fixation. Sd extends infection persistency under complete CI suppression.

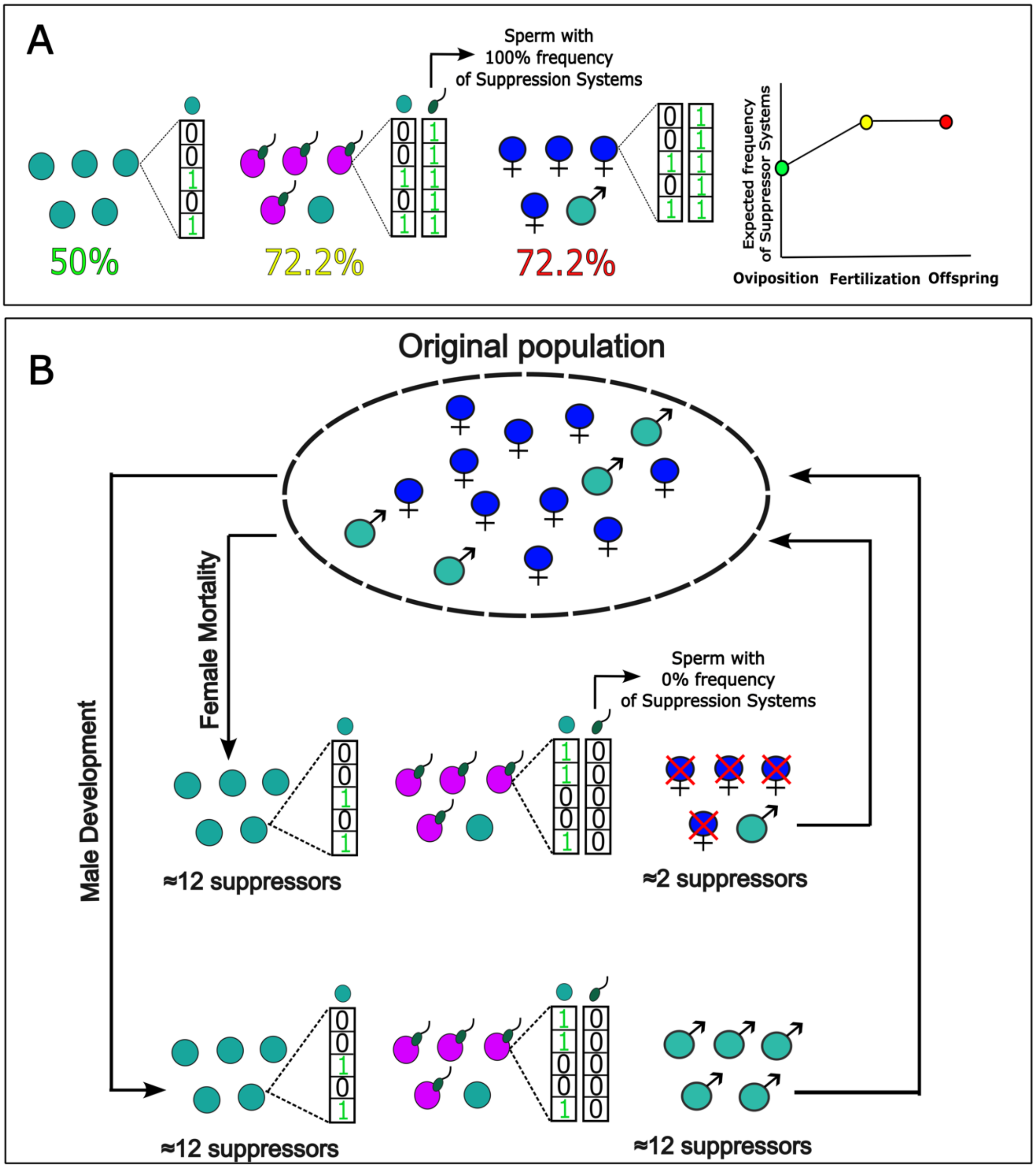

**Figure S7. Differential effects of CI phenotypes on the recycling of suppressors in a haplodiploid population.** The diagram shows the two types of CI crosses in haplodiploids, i.e., MD-CI and FM-CI. For males devoid of suppressors, CI strength is complete. For an expected number of ~12 suppressor alleles (50% frequency of host suppressors in the population) among offspring under both CI types, FM-CI erases suppressors from offspring, whereas MD-CI conserves these alleles.

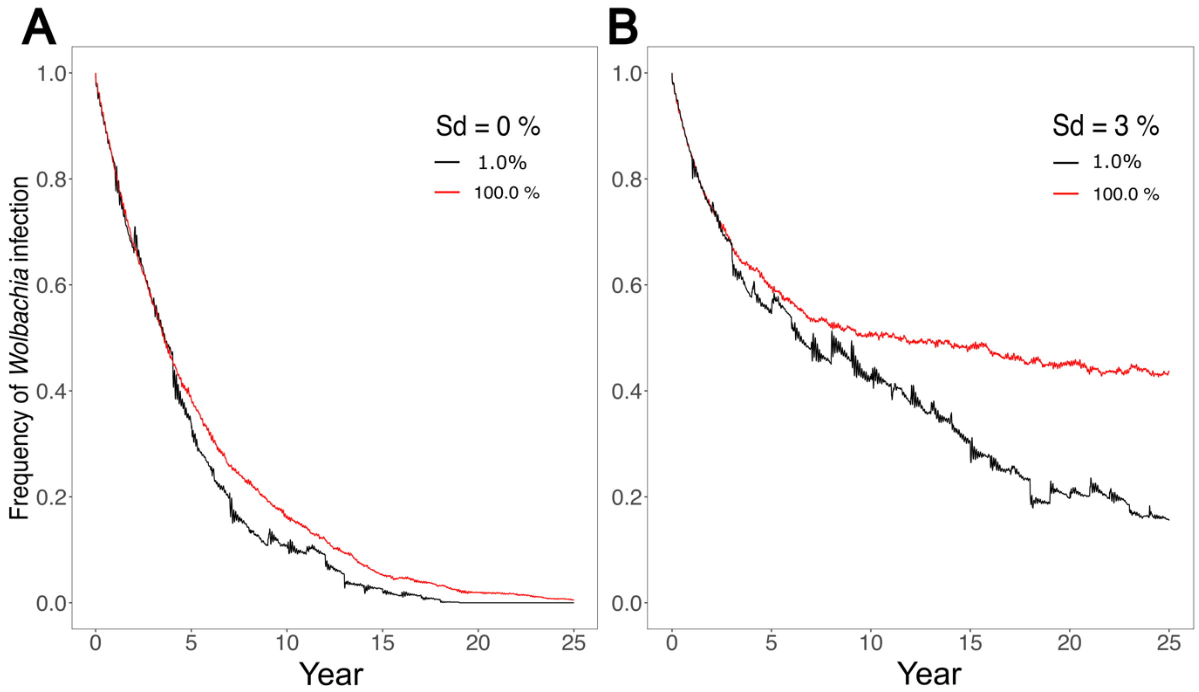

**Figure S8. Effect of drift on infection persistency.** In panels A and B, we show the effect of drift on infection frequencies across years. Drift is measured as the proportion of a population at the end of a given year that is carried over as the starting population of the next reproductive year. These proportions are given in the inset, with 1% representing approximately 50 individuals (for K = 5000). The effect of drift is particularly pronounced for a population under Sd, where infection persistency is extended.

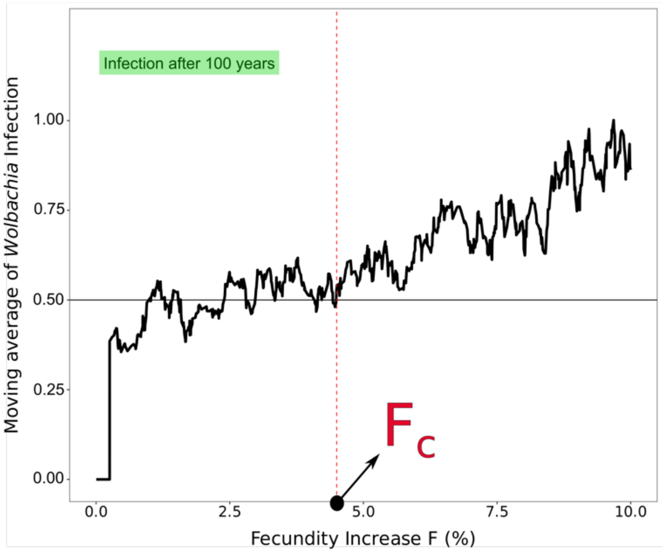

**Figure S9. Moving average of infection frequency as a function of *Wolbachia*-induced fecundity increase with a period of 10 observations.** The figure depicts an example of how a threshold value of $F$ ($F_c$) is calculated. For each fecundity value ($F$) we ran our simulation for a total of 100 years, with 100 independent runs, and retrieved the final infection frequency. We subsequently ordered the average infection frequencies according to an increasing sequence of $F$, after which we computed the moving average with a lag period of ten observations. In the illustrative example, the threshold value of fecundity ($F_c$) is the fecundity value for which the moving average hits the value of the starting infection frequency (50%) and stays above this threshold for all $F > F_c$.

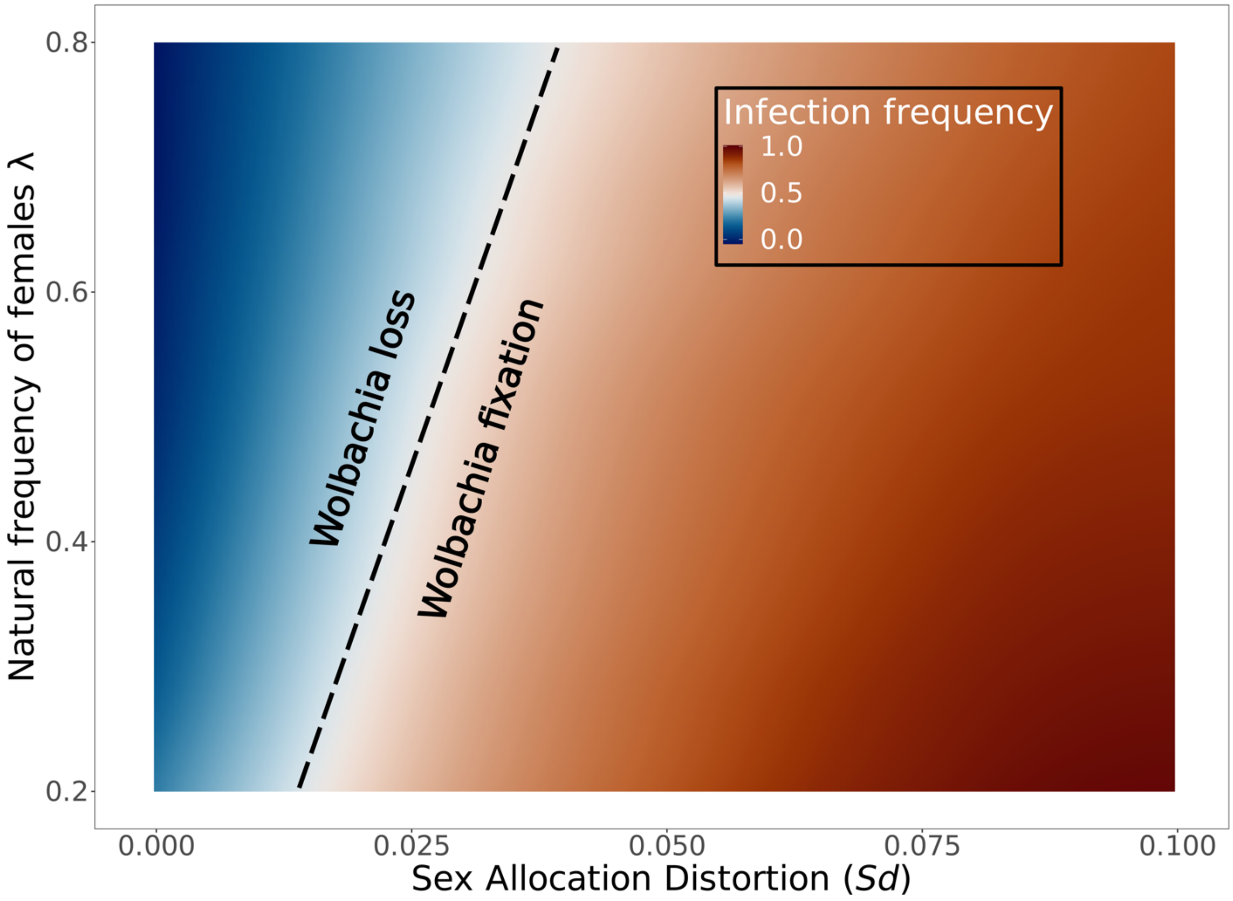

**Figure S10. Joint effects of the natural population sex ratio and sex allocation distortion on infection persistency.** Simulations with an initial infection frequency of 50% were performed for a total of 50 years. Final infection frequencies are depicted. If the final infection frequency is the same as the initial infection frequency, we verified that the symbiotic association was stable (indicated by the dashed black line), with deviations in the final infection frequency associated with loss or fixation of *Wolbachia*. The effect of Sd changes as a function to the natural population sex ratio. For high values of male to female ratios (low $\lambda$), *Wolbachia*-induced Sd has a much stronger effect on infection persistency.
